# CD8^+^ T cells regulate the bioenergetic reprogramming of lymphoid organs and the heart during viral infection

**DOI:** 10.1101/2025.08.20.670307

**Authors:** Laura Antonio-Herrera, Cecile Philippe, Attila Kiss, Hatoon Baazim, Juan Sanchez, Henrique Colaço, Felix Clemens Richter, Joel Xu En Wong, Fabian Amman, Magdalena Siller, Hao Wu, Sophia M. Hochrein, Stefanie Marie Ponti, Victoria Weissenböck, Thomas Wanek, Lukas Weber, Christopher Dostal, Seth Hallström, Elisabeth De Leeuw, Cedric Bosteels, Judith Lang, Usevalad Ustsinau, Xiang Li, Ferdinand Seith, Barbara F. Schörg, Anna Würth, Martina Schweiger, Philipp Starkl, Aubrey Burret, Anna Hofmann, Bethany Dearlove, Csilla Viczenczova, Alexander Lercher, Jakob-Wendelin Genger, Clarissa Campbell, Thomas Scherer, Anna Orlova, Richard Moriggl, Adelina Qerimi, Johannes Haybaeck, Rudolf Zechner, Sylvia Knapp, Karl S. Lang, Manfred Kneilling, Stefan Kubicek, Martin Vaeth, Bart N. Lambrecht, Bruno Podesser, Marcus Hacker, Andreas Bergthaler

## Abstract

The activation of the immune system is a bioenergetically-costly process^1^. Yet, essential bodily functions require a continuous energy supply, imposing energy constraints and trade-offs between competing processes^2^. Our understanding of the underlying bioenergetic adaptations reconciling rapid immune activation with other vital processes remains scarce. ^3–6^ Here, by using experimental models of viral infections, we identified an unexpected CD8^+^ T cell-driven redistribution of energy substrates between lymphoid organs and the heart. Viral infection promoted systemic hypoglycaemia and ketogenesis, together with systemic reallocation of energy substrates. Across organs analysed, secondary lymphoid organs and the heart showed the most dramatic changes. The former increased glucose uptake and oxidation while the heart showed the opposite, switching to preferential fatty acid utilization. These bioenergetic adaptations were absent in infected mice lacking CD8^+^ T cells or with T cells lacking the glucose transporter GLUT1. Pharmacological inhibition of fatty acid oxidation forced a systemic switch to glucose oxidation. This was associated with metabolic decompensation, reduced cardiac energetics, left ventricular stress, and mortality in otherwise nonlethal viral infections. Our results reveal how the energetic cost of immune cell activation imposes bioenergetic adaptations on non-lymphoid organs, posing a major challenge for the heart by completely relying on fatty acids.

## INTRODUCTION

Infections induce conserved sickness behaviours including anorexia^7,8^. This can affect energy availability, which under extreme circumstances may compromise survival^9^. The energy requirements of the pathogen, the activation of the immune system and the maintenance of essential bodily functions, can lead to a “war for glucose”^10^. Thus, paradoxically, the sick organism reduces food intake in periods of increased energy needs^7,8^.

The activation of adaptive immune cells is estimated to require an energy equivalent of up to 30% of the basal metabolic rate of the host, satisfied mostly from glucose utilization^1,11^. Positron emission tomography (PET) imaging of glucose shows increased uptake in secondary lymphoid organs (SLO) of patients with autoimmune disease, infections or upon vaccination^12–14^, suggesting increased glucose allocation to fuel immune activation. Still, how the organism re-allocates nutrients during periods of increased energy needs is poorly understood.

Organisms evolved mechanisms to preserve vital processes under stress conditions such as prolonged undernutrition and infections^15–18^. Life history of organismal biology indicates that increasing the energy expenditure of one biological process will deprive another one of energy if energy is limited^2^. These mechanisms ensure that energy resources are used for processes required for immediate survival. For instance, fueling of the immune system can compromise growth, reproduction and thermoregulation^15–17^. It stands to reason that failure to adequately re-allocate energy resources during extreme conditions such as infections will have a negative impact on host fitness. The exact biological processes, tissues, cells, and signals involved in energy trade-offs during infection remain under-characterized. In this study, we aimed to answer the fundamental question of how an infection-triggered immune response impacts organismal energy distribution, and how the bioenergetic coordination between immune and non-immunological organs supports host fitness.

## RESULTS

### Viral infections promote systemic changes in the abundance of energy substrates

First, we investigated whether viral infections affect systemic levels of energetic substrates, which would be indicative of differential mobilization or utilization. To account for host bioenergetic adaptations tailored to different pathogens, we analysed circulating glucose and ketone bodies (KB) during the course of infection. We studied two models of systemic viral infection with either the lymphocytic choriomeningitis virus (LCMV) strain Cl13 (chronic infection) or LCMV strain Armstrong (acute infection) ^19–21^, as well as two models of localized pulmonary infection with pneumonia virus of mice (PVM) strain J3666 ^22^ and with influenza A strain PR8/34 ^23^. The dynamic changes in food intake, body weight and blood metabolites were specific to each infection model (**Fig. 1a-c** and **Figure S1**). Infection with LCMV Armstrong promoted a mild reduction in food intake, body weight and hypoglycaemia but not ketogenesis (**Figure S1a**). Infection with influenza virus resulted in a pronounced drop in food intake and body weight but did not promote changes in blood glucose and KB (**Figure S1b**). In contrast, infection with LCMV Cl13 promoted reduced food intake, body weight loss and had the biggest impact lowering glucose levels and promoting increased KB levels in the blood (**Fig. 1c**). Together with an elevation of long-chain acylcarnitines levels in the serum (**Fig. 1d**), these data indicated mobilization of fat stores with subsequent engagement of fatty acid oxidation (FAO). PVM infection led to changes that were comparable to LCMV Cl13 infection regarding glucose and KB (**Extended Data Fig. 1c**). Moreover, PVM infection led to elevated serum levels of non-esterified fatty acids (**Figure S1d**), similar to LCMV Cl13 infection^24^. The profound changes in the abundance of energy substrates observed in these last two models of infection prompted us to investigate their corresponding adaptations in systemic energy utilization.

**Fig. 1:**
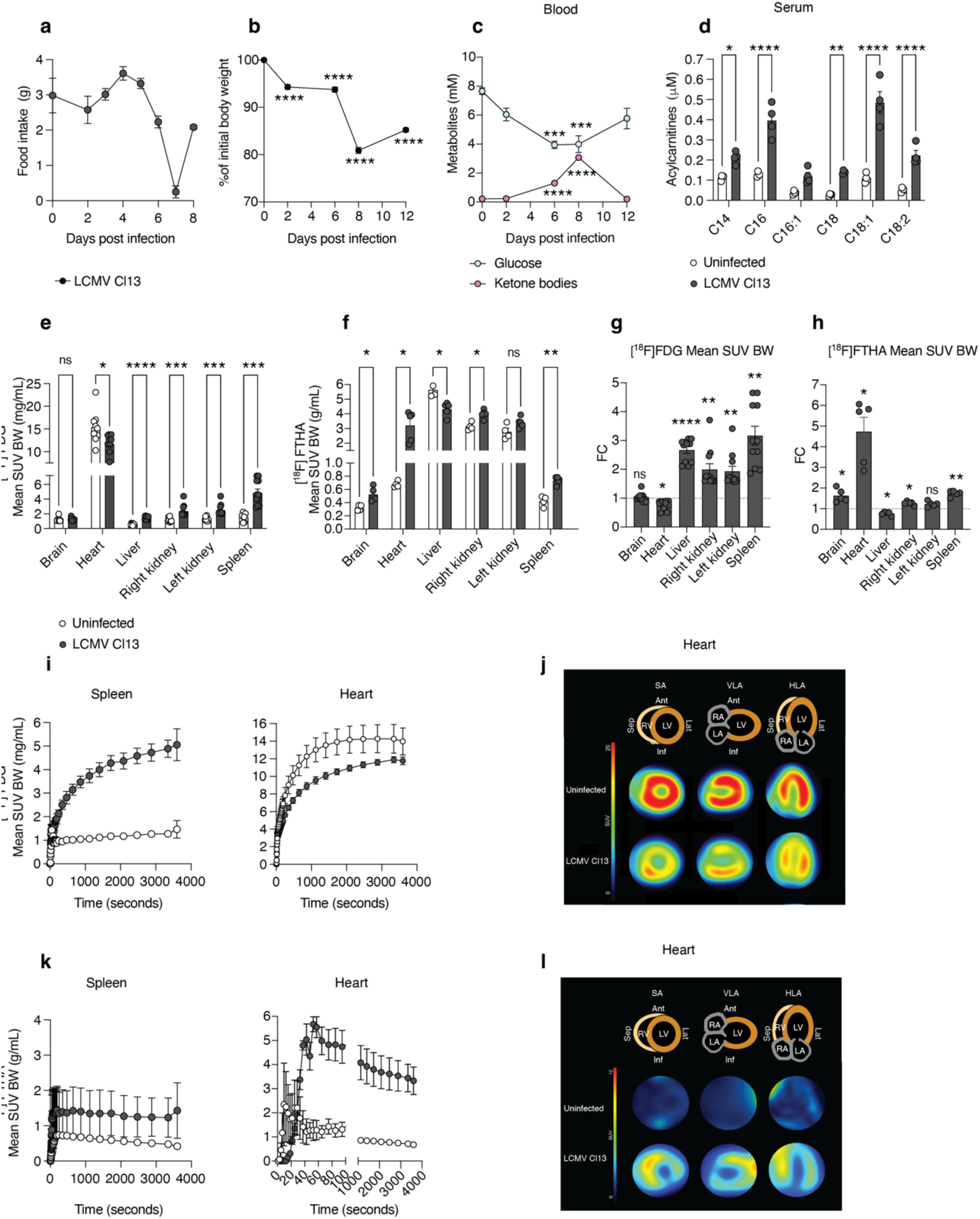
Energy constraints and reallocation during the adaptive immune phase of viral infection. a-d, WT mice were infected intravenously with 2 x 10^6^ FFU of LCMV Cl13, food intake and body weight were recorded and blood samples were taken at different time points for measurement of glucose, ketone bodies and acylcarnitines (the latter only at day 0 and 8). N= 5 mice per group. Statistical significance against their respective time zero (a). **e,** After eight days mice received an i.v. bolus of 2-deoxy-2-[^18^F]fluoro-D-deoxyglucose [^18^F]FDG or **f**, 14(R,S)-[^18^F]fluoro-6-thia-heptadecanoic acid [^18^F]FTHA and were scanned for 60 min by PET/CT. Mean standardized uptake values normalized to body weight (SUV BW) across organs after 60 min. **e**, N=9-11 mice per group, data pooled from three independent experiments. **f**, N=5 mice per groups, data pooled from two independent experiments. **g** and **h** depict the fold change (FC) of FDG and FTHA uptake values from **e** and **f**, respectively. **i**, Time activity curves showing SUV BW values during the whole PET/CT scan of spleen and heart, respectively. **j**, Representative images of [^18^F]FDG uptake in myocardial slices after 60 min. The top panel depicts the distinct areas of the heart visualized in every panel below them. SA, short axis, HLA, horizontal long axis, VLA, vertical long axis. RV right ventricle, LV left ventricle, RA right atrium, LA left atrium. **k**, Time activity curves of [^18^F]FTHA uptake in the heart and spleen. N=5 per group. **l**, Representative images of [^18^F]FTHA uptake in myocardial slices after 60 min. Unpaired T test, \**p* <0.05, \*\**p* <0.005, \*\*\**p* <0.0005, \*\*\*\**p* <0.0001.

### Redistribution of glucose and fatty acids between lymphoid and non-lymphoid organs during viral infection

We next aimed at characterizing the changes in energy distribution during viral infection. To obtain unbiased insights into the uptake of energy substrates at the organismal level, we employed whole-body positron emission tomography coupled to computer tomography (PET/CT) to track the uptake of the glucose analogue 2-deoxy-2-[^18^F]fluoro-D-deoxyglucose ([^18^F]FDG) in mice infected with LCMV Cl13 and compared them to uninfected mice. This was done eight days after the inoculation of the virus, when the most dramatic changes in blood glucose and KB were observed (**Fig. 1c**). Glucose uptake was enhanced in various organs of infected mice including liver, kidneys and in SLO such as the spleen and lymph nodes (**Fig. 1e** and **Figure S2a-c**). No significant changes were observed in skeletal muscle or brain (**Fig. 1d** and **Figure S2c**).

Having observed an enrichment of KB in the blood, indicating activation of FAO upon LCMV Cl13 infection, we next aimed at dissecting fatty acid uptake at the systemic level. We administered radioactively labelled long-chain fatty acid 14(*R,S*)-[^18^F]fluoro-6-thia-heptadecanoic acid ([^18^F]FTHA) and observed infection-dependent changes in fatty acid uptake in most quantified organs except for lymph nodes and skeletal muscle (**Fig. 1f** and **Figure S2d-f**).

Of note, the spleen showed one of the highest increases in glucose uptake (**Fig 1g, i**), whereas the heart was the only organ with a significant reduction, specifically in the myocardium and excluding the blood-filled ventricles and atria (**Fig. 1g,i,j**).

These data indicate that during LCMV Cl13 infection multiple organs, lymphoid and non-lymphoid, maintain or increase their uptake of glucose with the exception of the heart. At the same time, the most dramatic change was the enhanced uptake of FTHA in the myocardium of LCMV Cl13 - infected mice compared to uninfected controls (**Fig. 1h, k,l**). These data revealed discrete changes in fatty acid utilization across organs during infection, with a considerable allocation of fatty acids to the heart.

### Glucose supports glycolysis in the spleen whereas fatty acids support energy metabolism in the heart during viral infection

Having observed a differential uptake of glucose and fatty acids during the adaptive phase of the immune response to a viral infection, we focused next in the utilization of these metabolites for energy production in spleen and heart. We first dissected the incorporation of heavy isotope labelled ^13^C_6_-glucose into the main energy pathways glycolysis and the tricarboxylic acid cycle (TCA) (**Fig. 2a**). In the spleens of infected mice, glucose contribution to the TCA was less compared to uninfected animals (**Fig. 2b**), while the tracer was highly enriched in the glycolytic products pyruvate and lactate (**Fig. 2c**). This suggests that glucose contributes to energy production mainly through anaerobic glycolysis in the spleen at this time of the infection. Of note, the total pool of the TCA and glycolytic intermediates were increased or remained unaffected in infected mice (**Figure S3a,b**). In the hearts of infected mice, our tracing data showed that glucose contribution to both the TCA and glycolysis was severely dampened (**Fig. 2d,e**). The concentrations of the total pool of TCA metabolites and glycolysis products pyruvate and lactate was, however, not compromised in heart tissue of infected mice compared to uninfected controls (**Figure S3c,d**). Despite reduced glucose uptake and utilization, this indicates that the total production of energy metabolites in the heart is not impaired in infected mice. Considering our PET/CT imaging data, this hinted at compensation by other metabolites such as fatty acids.

**Fig. 2:**
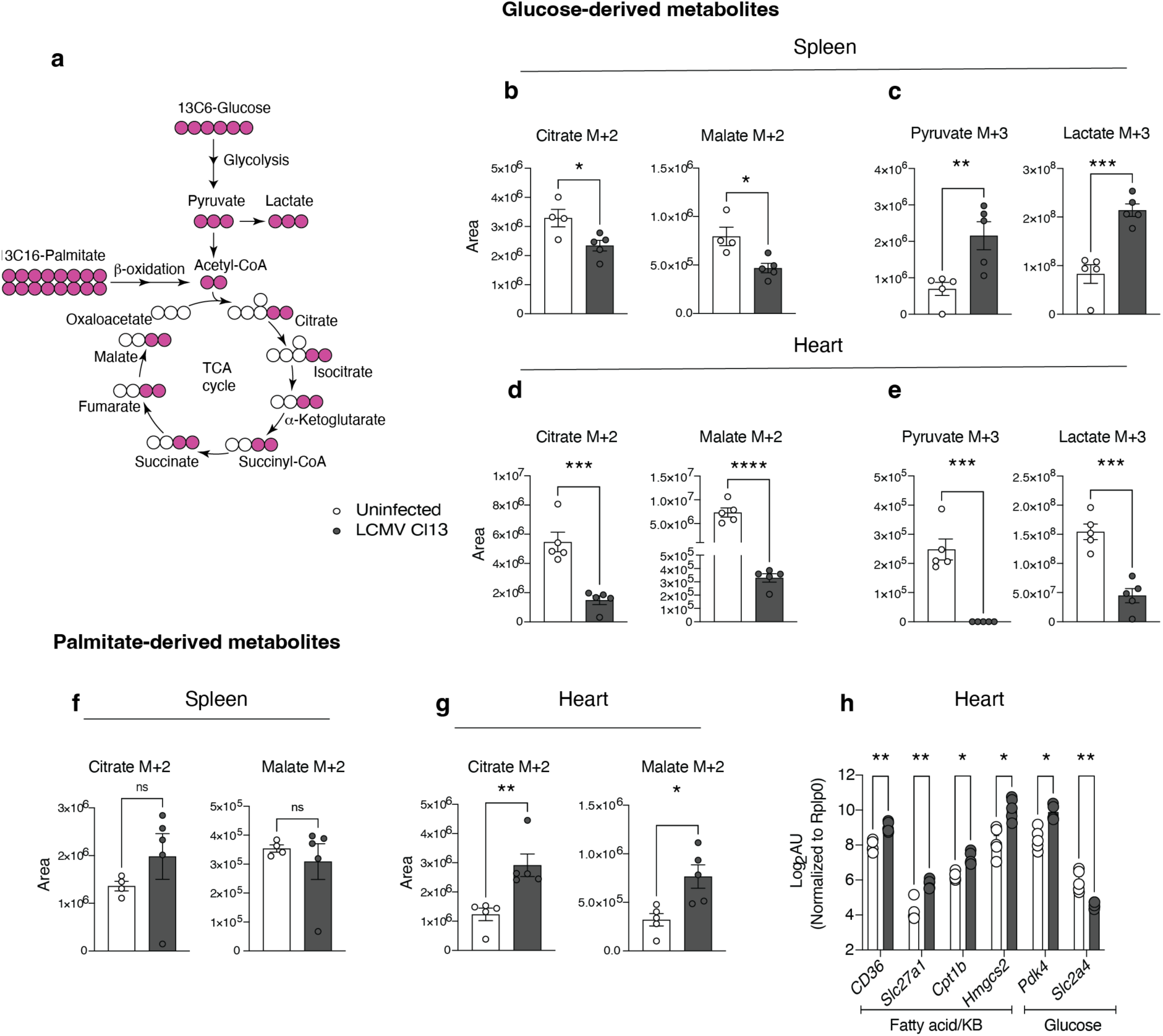
Spleen glucose sparing and pronounced fatty acid oxidation in the heart characterize the adaptive immune phase of a viral infection. **a**, Diagram summarizing glycolysis and the TCA cycle, as well as the incorporation of ^13^C derived from either glucose or palmitate. Mice were infected with LCMV Cl13, after eight days they received an i.p. injection of ^13^C_6_-glucose (**b**-**e**) or ^13^C_16_-palmitate/BSA (**f, g**), tissues were recovered after 10 or 20 min., respectively and analysed by mass spectrometry, the data is represented as area under the curve. **b**, **d** Shows ^13^C-labeled TCA metabolites in the spleen and heart and **c**, **e** shows ^13^C-labeled glycolysis products in the spleen and heart upon the injection of ^13^C_6_-glucose. **f**, **g**, Shows ^13^C-labeled TCA metabolites in the spleen and heart, respectively. N=4-5 mice per group, the data is representative of three (b-e) and two (f, g) independent experiments. **h**, Heart expression of genes involved in fatty acid and glucose metabolism in uninfected mice or mice infected for eight days. N=5 mice per group, data is representative of at least two independent experiments. Data is showed as the mean ± s.e.m. Unpaired T test, \**p* <0.05, \*\**p* <0.005, \*\*\**p* <0.0005, \*\*\*\**p* <0.0001.

To study the contribution of fatty acids to energy production, we interrogated their incorporation into the TCA by administering a single dose of heavy isotope labelled ^13^C_16_- palmitate on day 8 after infection (**Fig. 2a**). The incorporation of the tracer in spleen was comparable between infected and uninfected samples (**Fig. 2f**). In the heart tissue of infected mice, ^13^C incorporation was significantly enriched in the TCA metabolites compared to uninfected controls (**Fig. 2g**). Together, these results corroborate that glucose increasingly contributes to energy production in the spleen during viral infection while circulating fatty acids have a greater contribution in supporting energy production in the heart.

To further understand the bioenergetic adaptations taking place in the heart during viral infection, we analysed the expression of genes relevant for fatty acid and glucose utilization. We found increased expression of genes involved in fatty acid transport (*Cd36*, *Slc27a1*), FAO (*Cpt1b*), and KB synthesis (*Hmgcs2*) upon infection with LCMV Cl13 (**Fig. 2h**). The increased expression of *Pdk4*, a negative regulator of glucose utilization that facilitates FAO^25^ together with decreased expression of *Slc2a4* (**Fig. 2h**), coding for the main heart glucose transporter GLUT4, further supported that the heart engages into preferential fatty acid utilization. These transcriptomic changes were partially reproduced in the unrelated infection model of PVM, whereby, next to hypoglycaemia and ketogenesis, we also observed the upregulation of *Slc27a1*, *Hmgcs2 and Pdk4* as well as a trend to lower *Slc2a4* expression in the heart (**Figure S3e**). Thus, infections promoting systemic hypoglycaemia and ketogenesis may also trigger bioenergetic adaptations in the heart leading to preferential fatty acid utilization. Together, our results show that the organism can adapt to viral infections through the reallocation and differential utilization of energy substrates across SLO and vital organs like the heart.

### CD8^+^ T cells regulate the abundance and redistribution of energy substrates in spleen and heart during viral infection

We next aimed to identify the mediator(s) of the energetic adaptations seen upon viral infection. First, we assessed the impact of reduced food intake on the availability of circulating energy substrates. Uninfected mice received the same amount of food that mice infected with LCMV Cl13 consume daily during eight days. This pair-fed uninfected group showed non-significant changes in either blood glucose or KB (**Fig. 3a**), suggesting that reduced food intake in the LCMV Cl13 infection model is not sufficient to induce altered blood levels of glucose or KB. Recently, insulinemia was found to correlate with blood glucose during the innate phase of the antiviral immune response against LCMV Cl13 ^26^. Having observed that the most pronounced changes in both glucose and KB take place during the adaptive phase, we measured serum insulin at eight days post infection but found no differences compared against uninfected mice (**Fig. 3b**), arguing against a dominant role for insulin at this later time point of infection.

**Fig. 3:**
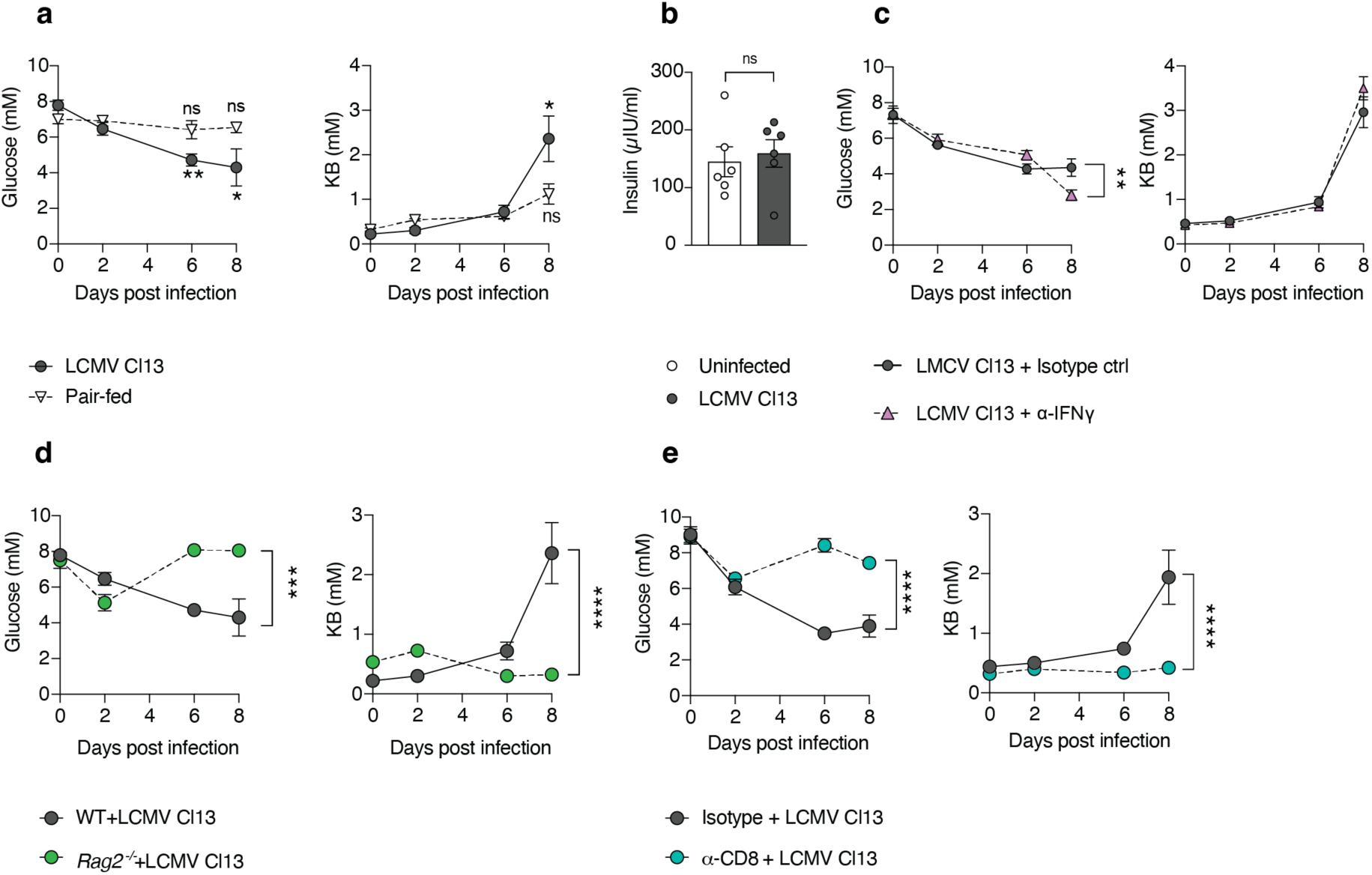
CD8 T cells regulate the concentration of circulating glucose and ketone bodies during the adaptive immune phase of viral infection. **a**, Food intake of mice infected with LCMV Cl13. Body weight loss of LCMV Cl13 uninfected mice received the average amount of food consumed by LCMV Cl13 infected mice daily (pair fed). Blood glucose and KB were measured at the indicated time points. N=5, the data is representative of two independent experiments. Statistical significance against their respective time zero. **b**, mice were infected with LCMV Cl13 and serum was obtained after eight days to measure insulin. N= 6 mice per group, data pooled from two independent experiments. **c**, Mice were treated with an anti-IFNγ antibody, or an isotype control, before and during the infection with LCMV cl13. Blood glucose and KB were measured at the indicated time points. N=8-10, data pooled from two independent experiments. Statistical significance against their respective time zero. **d**, WT or *Rag2*^-/-^ mice were infected with LCMV Cl13 and blood glucose and KB were measured at various time points, N= 4 mice per group. Data is representative from two independent experiments. **e**, Mice were treated with a blocking antibody against CD8 or an isotype control before LCMV Cl13 infection. The graphs show glucose and KB concentrations in the blood at different time points. N= 5 mice per group. Data is representative from two independent experiments. Statistical significance against their respective time zero. Data is showed as the mean ± s.e.m. Unpaired T test. \**p* <0.05, \*\**p* <0.005, \*\*\**p* <0.0005, \*\*\*\**p* <0.0001.

A recent study by Šestan^26^ et al. also showed how IFNγ signalling can regulate blood glucose during the early phase of viral infection. When we neutralized IFNγ by administration of a blocking antibody at eight days post infection, we observed enhanced infection-induced hypoglycaemia and no change in blood KB compared to control (**Fig. 3c**). This suggested that IFNγ is not the primary promoter of LCMV Cl13-induced hypoglycaemia and ketonemia in the adaptive immune phase.

The results made us hypothesize that the mechanisms regulating the abundance of energy metabolites during the adaptive immune response against the infection are different to those during the innate phase. We previously demonstrated that T cell responses mediate profound systemic metabolic changes during LCMV Cl13 infection, including body weight loss, loss of adipose and skeletal muscle mass, as well as increased levels of circulating fatty acids^24^. Thus, we decided to investigate the role of T cell response by employing the *Rag2^-/-^*model which lacks mature lymphocytes. Upon infection, blood glucose and KB changes observed after six-eight days post infection in WT mice were absent in infected *Rag2^-/-^* mice (**Fig. 3d**). The same outcome was achieved upon antibody-mediated depletion of CD8^+^ T cells (**Figure S4a, Fig. 3e**). Similar observations were made in the model of PVM infection as *Rag2^-/-^*mice were protected against infection-induced body weight loss, hypoglycaemia and ketogenesis (**Figure S4b,d,e**). Ablation of CD8^+^ T cells resulted in comparable trends in PVM-infected WT animals (**Figure S4c-e**). Overall, our data indicate that T cells regulate the abundance of energy metabolites during the adaptive immune response against a viral infection.

We next investigated whether T cells mediate also the infection-induced bioenergetic reprograming of spleen and heart. For this we performed PET/CT imaging studies whereby animals received a bolus of the glucose analogue 2-DG or the long-chain fatty acid FTHA coupled to radioactive tracer. Infected mice lacking CD8^+^ T cells showed reduced uptake of glucose in the spleen and an inverse patter in the heart (**Fig. 4a**). These data implied that CD8^+^ T cell responses are necessary for the observed increased glucose demands of SLO and decreased glucose uptake in the heart upon viral infection.

**Fig. 4:**
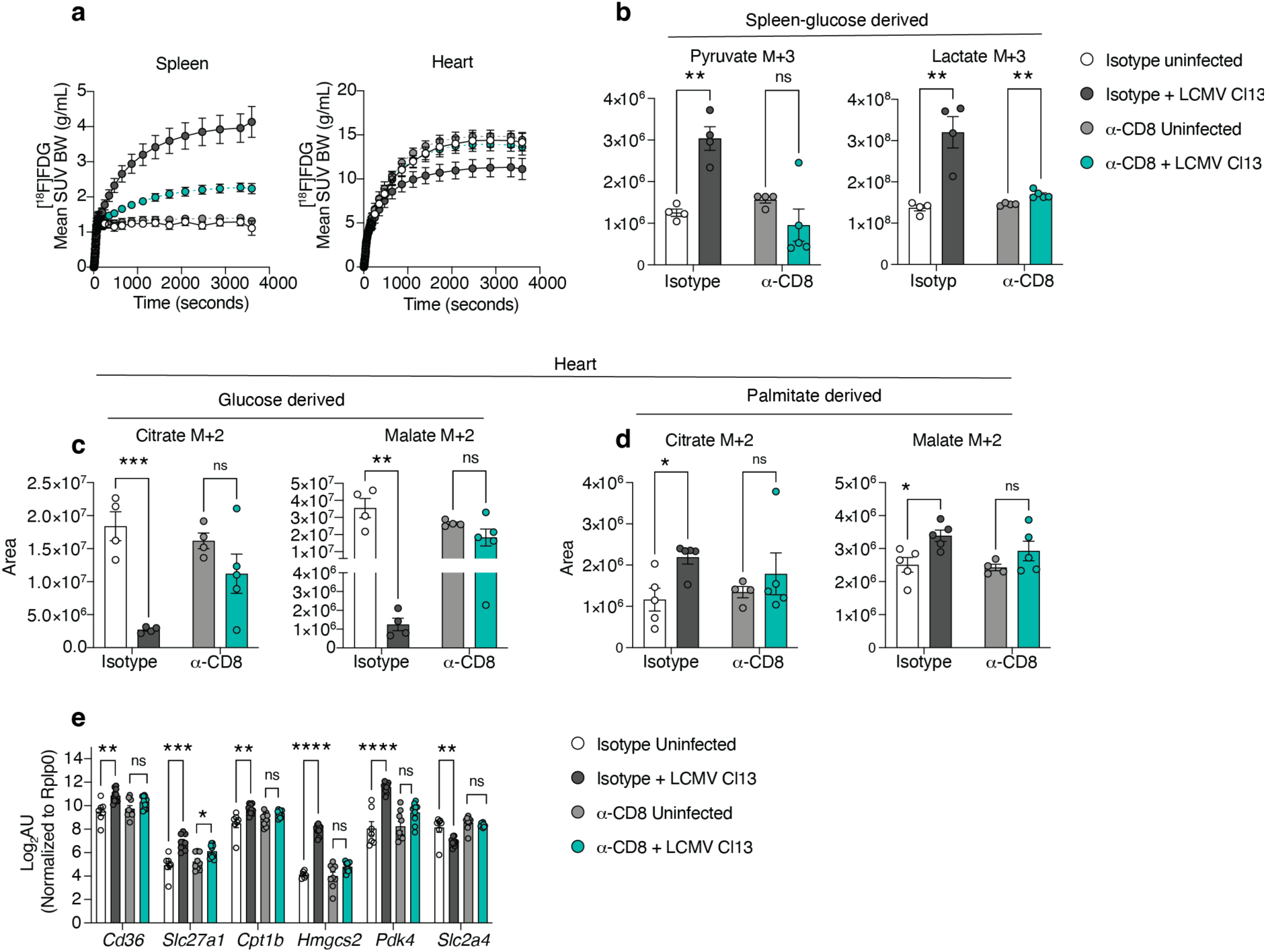
CD8 T cells promote the shifts in glucose and fatty acid utilization in the heart. **a**, Mice were treated with a blocking antibody against CD8 or an isotype control before LCMV Cl13 infection. After eight days mice received one injection of [^18^F]FDG and tracer uptake was analysed by PET/CT for 60 min. The graphs show time activity curves of tracer uptake in spleen and heart. N= 6-8 mice per group. Data poled from two independent experiments. **b-d**, Alternatively, mice received one injection of ^13^C_6_-glucose or ^13^C_16_-palmitate eight days after the infection. 10 or 20 minutes later, respectively, organs were recovered for mass spectrometry analysis. N=4-5 mice per group. e, Gene expression in the heart eight days after infection. N=9-10 mice per group. Data was pooled from two independent experiments. The data represent the mean ± s.e.m. Unpaired T test. **g**, \**p* <0.05, \*\**p* <0.005, \*\*\**p* <0.0005, \*\*\*\**p* <0.0001.

Yet, it remained unclear whether CD8^+^ T cell depletion is sufficient to prevent the changes in energy substrate utilization. To address this, we depleted mice of CD8^+^ T cells and administered a single dose of stable isotype ^13^C_6_-glucose to measure its incorporation into the TCA and glycolysis pathways and the total pool of these metabolites (**Fig. 2a**). In the spleen and heart, the concentrations of the pool of TCA and glycolysis metabolites were comparable between uninfected and infected mice upon anti-CD8 antibody treatment (**Figure S4f,g**). In the spleen, viral infection promoted increased utilization of glucose to fuel anaerobic glycolysis (**Fig. 2c**). CD8^+^ T cell depletion prevented the latter, resulting in similar glucose incorporation into glycolysis between uninfected and infected animals, unlike the isotype treated controls (**Fig. 4b**). These results indicate that anaerobic glycolysis in the spleen was linked to the presence of CD8^+^ T cells. As shown before (**Fig. 2d,e**), viral infection resulted in a reduced contribution of glucose to anaerobic glycolysis and TCA cycle in the heart. Again, CD8^+^ depletion prevented this phenotype, resulting in similar incorporation of glucose into the TCA in the hearts of infected and uninfected mice (**Fig. 4c**). This indicates that antiviral CD8^+^ T cell responses impair glucose utilization by the heart for energy production via the TCA cycle. Next, we investigated whether CD8^+^ T cells drive the cardiac utilization of fatty acids promoted by LCMV CL13 infection (**Fig. 2g**). Indeed, CD8^+^ T cell depletion prevented the increased incorporation of ^13^C_16_-palmitate into TCA intermediates (**Fig. 4d**). Next to the lack of infection-induced adaptations in energy substrate utilization, we did not observe neither the transcriptional changes in the heart linked to glucose and fatty acid utilization upon CD8^+^ T cell depletion (**Fig. 4e**). Similarly, the upregulation of *Hmgcs2* and *Pdk4* in heart tissue of PVM-infected mice was not observed in the hearts of *Rag2*^-/-^ or CD8^+^ T cell depleted mice (**Figure S4h,i**). Together our data indicate that adaptions in energy substrate utilization during viral infection are mediated by T cells.

Of note, mice lacking CD8^+^ T cells had higher viral loads in blood and expression of viral protein in the heart than isotype-treated infected animals (**Figure S4j,k**). This confirms the key role of CD8^+^ T cells in antiviral control and, at the same time, argues against prolonged viral replication as critical mediator of the bioenergetic host adaptations.

Together, our data underline the impact of T cell activation on the abundance, redistribution and utilization of energy substrates during a viral infection.

### Impact of glucose consumption by T cells on systemic bioenergetic adaptations

Our results show that CD8^+^ T cells promote a switch in systemic energy preferences, impacting mostly the heart and SLOs. However, the question about how CD8^+^ T cells would mediate these systemic effects, and the involvement of key aspects of CD8^+^ T cell biology such as tissue migration, cytotoxicity, and cell-intrinsic energy demands, remained open.

To study the impact of CD8^+^ T cell migration, we administered infected mice with the drug FTY720, which inhibits the exit of lymphocytes from SLOs into the periphery^27^. FTY720 treatment reduced T cell egress from SLO, as suggested by the retention of CD8^+^ T cells in spleen and reduced numbers in the blood of infected mice (**Figure S6a**). This, however, had no impact on either blood glucose or KB levels during the infection (**Fig. 5a**), suggesting a negligible role for T cell migration to peripheral tissues to modulate energy substrate abundance. Next, we investigated the role of CD8^+^ T cell cytotoxicity by employing animals lacking the expression of the key cytotoxic effector perforin (*Prf1^-/-^*). Upon infection, these animals showed no difference in the blood levels of either glucose or KB, compared to infected WT controls (**Fig. 5b**). These data suggested that classical CD8^+^ T cell cytotoxicity does not have a major impact on the systemic abundance of energy substrates. Further supporting these observations, we did not find correlations between organ-specific viral loads and CD8^+^ T cell infiltration with glucose or fatty acid uptake (**Figure S5**). Together, our data indicate that local effects of CD8^+^ T cells, including viral control and cytotoxicity, are unlikely to be the main drivers of systemic bioenergetic adaptations.

**Fig. 5:**
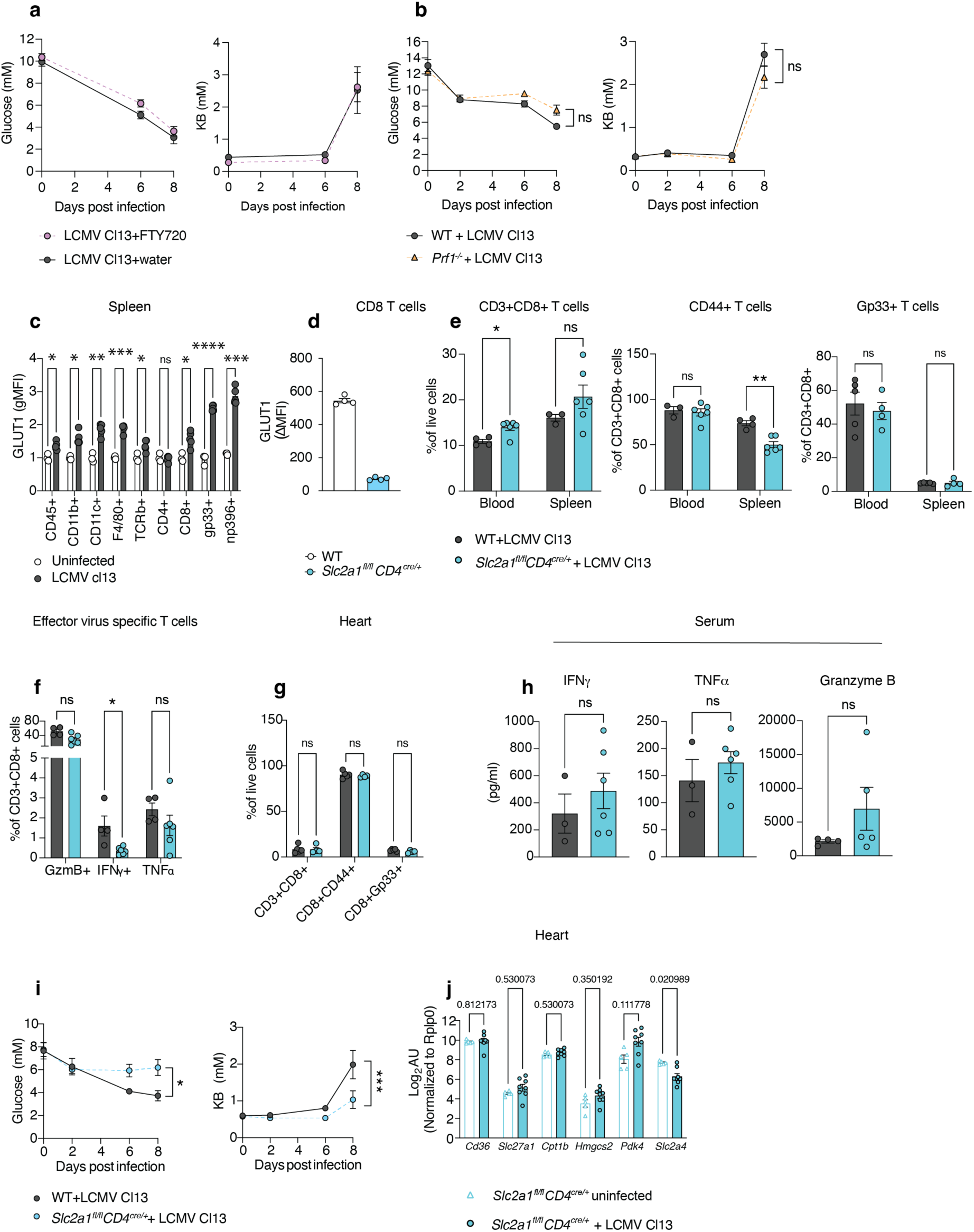
Glucose consumption by CD8 T cells partially regulates systemic abundance of energy metabolites and heart reprograming. WT mice were infected with LCMV Cl13. After four days they received an injection of FTY720 every day till day eight. **a**, Blood glucose and KB measured at the indicated time points. N=4 mice per group from one experiment. **b**, Blood glucose and KB of WT and *Prf1^-/-^* mice infected with LCMV Cl13. **c**, GLUT1 staining of splenocytes showing expression in various immune cell types. N=5 mice per group, data is representative of two independent experiments. **d**, GLUT1 staining of CD8^+^ T cells in vitro activated with α-CD3/CD28 antibodies. The data is expressed as median fluorescence intensity (MFI) subtracting the fluorescence minus one staining, N= 4 mice per group, data is representative of two independent experiments. **e**, WT and *Slc2a ^fl/fl^ CD4^cre/+^* mice were infected with LCMV Cl13, eight days later spleens and blood were recovered for FACS analysis of total CD8^+^, CD8^+^CD44^+^, Gp33^+^ and, **f**, cytokine producing T cells upon in vitro re-stimulation of splenocytes with Gp33 peptide. **g**, T cell infiltrates in the heart measured by FACS. N=4-6 mice per group. Data is representative of two independent experiments. **h**, Measurement of cytokines in serum. N=3-6 mice per group, data pooled from two independent experiments. **c-h**, unpaired T test. **i**, Blood glucose and KB were measured at the indicated time points. N= 8-9 mice per group, data pooled from two independent experiments. Two-way ANOVA. **j**, qPCRs from heart tissue. N=5-8 mice per group, data was pooled from two independent experiments. Unpaired T test. Data is showed as the mean ± s.e.m. \**p* <0.05, \*\**p* <0.005, \*\*\**p* <0.0005, \*\*\*\**p* <0.0001.

Next, we investigated whether glucose utilization by T cells, independent of effector function, influences systemic changes in energy utilization. Previously, the glucose receptor GLUT1 encoded by *Slc2a1* was shown to be essential for the activation of T cells^28^. When we assessed GLUT1 expression in the spleen upon LCMV Cl13 infection, we found increased GLUT1 levels in many immune cell populations and highest in virus-specific CD8^+^ T cells (**Fig. 5c**).

Hence, we studied a genetic mouse model where T cells lack GLUT1 expression (*Slc2a1^fl/fl^ CD4^cre/+^*)(**Fig. 5d**) ^28^. CD8^+^ T cells from these mice activated *in vitro* for 24 h showed reduced glucose uptake (**Figure S6b**) and glycolysis but normal mitochondrial oxidative phosphorylation relative to WT cells (**Figure S6c-e**), demonstrating impaired energy production from glucose. *In vivo*, the lack of GLUT1 expression had a moderate impact on T cell activation eight days upon viral infection (**Fig. 5e-f**). *Slc2a1^fl/fl^ CD4^cre/+^* mice infected with LCMV CL13 showed no differences in the proportion of splenic CD8^+^ T cells (**Fig 5e**), but reduced numbers of CD44+ and IFNγ^+^ CD8^+^ T cells (**Fig. 5e, f).** Gp33+, Granzyme B^+^ and TNFα^+^ CD8 T^+^ cells were unaffected (**Fig. 5e, f**). The infiltration CD8^+^ T cells in the heart was not affected either (**Fig. 5g**). Serum levels of IFNγ, TNFα and granzyme B were unaffected in *Slc2a1^fl/fl^ CD4^cre/+^* mice compared to infected littermate controls (**Fig. 5h)** Viral loads were comparable to those found in WT mice (**Figure S6f**). Together, our data indicates that the lack of GLUT1 expression in the T cell compartment has a minor effect on T cell activation without affecting viral clearance by day eight after infection. Likely, potential compensatory mechanisms by other glucose transporters expressed in T cells could explain the lack of a significant T cell phenotype.

Strikingly, while mice of either genotype showed an initial drop in blood glucose upon infection, only wildtype mice showed a further reduction in blood glucose towards eight days after infection (**Fig. 1b** and **Fig. 5i**). Likewise, we observed only moderately elevated levels of blood KB in *Slc2a1^fl/fl^ CD4^cre/+^* mice compared to controls upon infection (**Fig. 5i**). This indicates that T cell-intrinsic impairment of glucose utilization impacts the systemic levels of energy metabolites.

We next aimed to dissect whether reduced glucose uptake of T cells affected the bioenergetic reprograming of the heart during viral infection. The hearts of *Slc2a1^fl/fl^ CD4^cre/+^* animals infected with LCMV Cl13 showed not significant differences in the expression of genes involved in FAO including *Slc27a1*, *Hmgcs2, Pdk4 and Slc2a4* compared to the expression levels of uninfected hearts (**Fig. 5j**). Interestingly, the expression levels of *Slc2a4* remained low (**Fig. 5j**). Thus, glucose uptake by T cells was at least partially driving the bioenergetic adaptations of the heart promoted by viral infection.

### Disruption of fatty acid utilization during the adaptive immune phase compromises cardiac bioenergetics and reduces survival to infection

Having observed that T cell responses promoted elevated levels of blood KB and a biased reprograming in the heart towards FAO, we next investigated the relevance of FAO for the host’s fitness during viral infection by pharmacological FAO blockade with etomoxir^29^. Two days after LCMV Cl13 infection, the administration of etomoxir had no overt effect on clinical symptoms compared to vehicle treatment (**Fig. 6a**). Strikingly, when etomoxir was administered six- or eight-days after LCMV Cl13 infection, i.e. at the peak of bioenergetic changes (**Fig. 1c**), mice reached the humane endpoint four hours after drug treatment (**Fig. 6a, b**). Within this time frame, these mice showed metabolic decompensation characterized by severe hypoketotic hypoglycaemia (**Fig. 6c**), and increased concentration of blood urea nitrogen (BUN) (**Figure S7a**). Markers for infection-associated liver damage alanine aminotransferase (ALT) and aspartate aminotransferase (AST), and the heart-damage marker creatine kinase-MB (CKMB) were comparable (**Figure S7a**). Notably, administration of the inhibitor of glucose utilization 2-deoxyglucose (2-DG) was not detrimental to host fitness in mice infected with LCMV Cl13 (**Fig. 6d**).

**Fig. 6:**
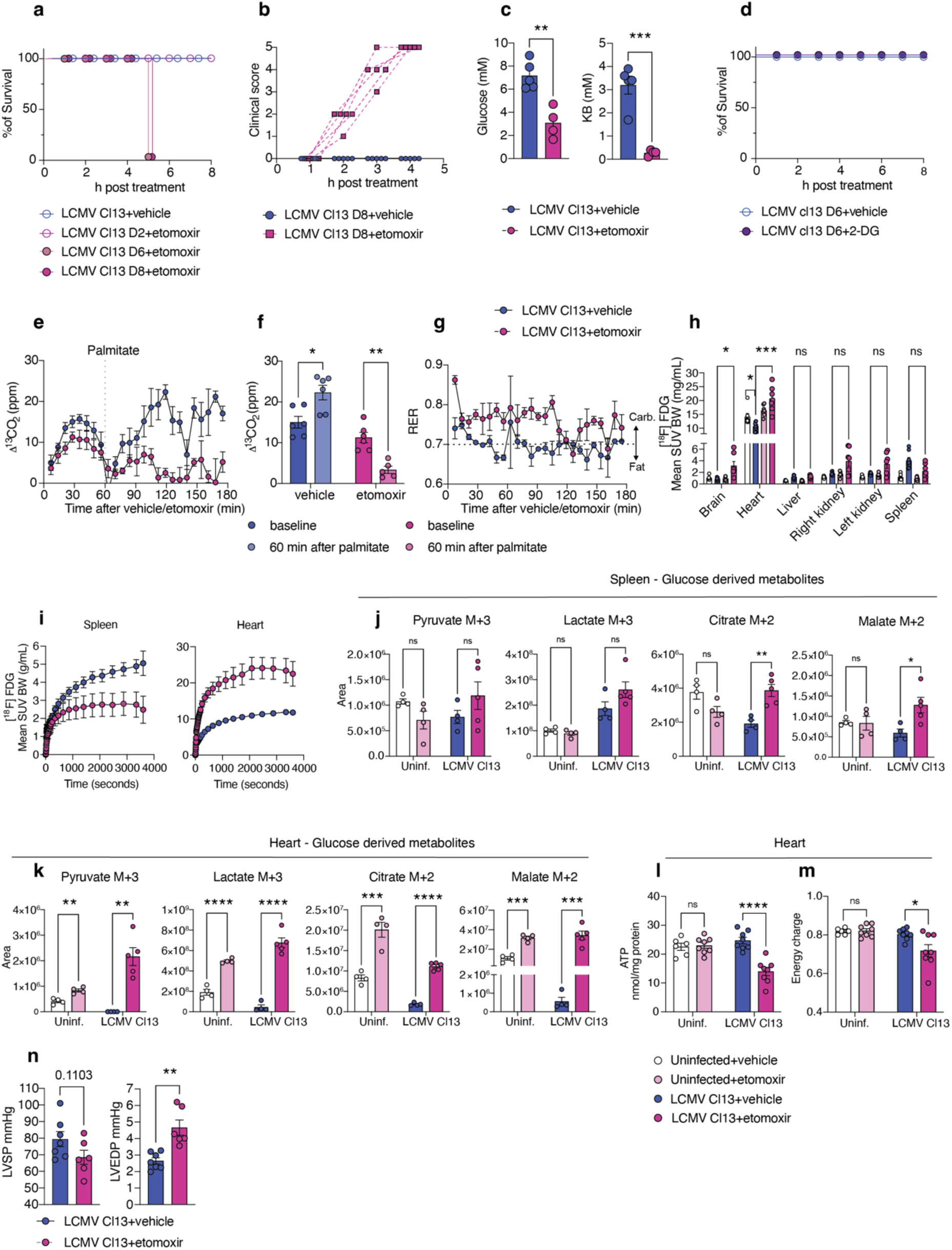
T cell induced metabolic reprograming supports heart bioenergetics and survival during viral infection. **a**, **b**, Mice infected with LCMV Cl13 received a single i.p. injection of etomoxir (20 mg/kg) or vehicle at the indicate time points. In **a**, 0 Indicates that mice reached the humane end point and were euthanized. N= 5 mice per group. The data is representative of at least two independent experiments. **c,** Glucose and KB were measured before and two hours after the injection of etomoxir N= 4-5 mice per group, the data is representative of at least two independent experiments. **d**, Mice were infected with LCMV Cl13 and after six days injected with 2-DG (500 mg/kg). **e**, Mice infected with LCMV Cl13 were treated with etomoxir after eight days and immediately housed in metabolic cages. One hour later, mice received one injection of ^13^C_16_-palmitate. The graph depicts the concentration of exhaled ^13^CO_2_ for three hours, **f**, shows the concentration after one hour of palmitate injection (two h after etomoxir). **g**, Respiratory exchange ratio (RER). n=6 mice per group, data pooled from two independent experiments. **h**, [^18^F]FDG uptake across organs two hours after etomoxir injection in mice previously infected with LCMV Cl13. The graph shows the mean SUV BW values at the end of the scans (60 min). **i**, Time activity curves of tracer uptake in spleen and heart. N= 7-8 animals per group, data pooled from two independent experiments. **j, k**, Mice treated with etomoxir or vehicle received one injection of ^13^C_16_-glucose after one hour. 20 minutes later organs were recovered for mass spectrometry analysis. The graphs show the ^13^C-labeling in TCA and glycolysis metabolites in the spleen and heart. N= 4-5 mice per group, the data is from one experiment. **l**, ATP normalized to protein content and **m**, energy charge of the heart two hours after vehicle or etomoxir injection. n= 6-8 mice per group, data pooled from four independent experiments. **n**, Invasive hemodynamics assessment two hours after drug treatment of mice infected with LCMV Cl13 eight days before. N= 5-7 mice per group, the data was pooled from two independent experiments. **l**, LVSP: left ventricular systolic pressure and LVEDP: left ventricular end diastolic pressure. Data is showed as the mean ± s.e.m. Unpaired T test. \**p* <0.05, \*\**p* <0.005, \*\*\*\**p* <0.0001.

We next aimed at investigating whether etomoxir injection could have a negative impact on mice in different contexts of reduced food intake, either in pair fed mice or mice infected with other viruses i.e., LCMV Armstrong, influenza and PVM. The lethal phenotype of etomoxir treatment was absent in uninfected mice fed the same amount of food consumed by LCMV Cl13-infected mice (**Figure S7b**), demonstrating that reduced food intake in the absence of infection is insufficient to render mice susceptible to lethal FAO blockade. Next, we administered a single dose of etomoxir to either mice infected with LCMV Armstrong or with Influenza PR8. The survival of these mice was not impaired following etomoxir injection (**Figure S7c,d**). Thus, infection alone is not sufficient to render mice susceptible to lethal FAO blockade.

We reasoned that the severity of FAO blockade with etomoxir manifests in models where the organism has shifted considerably towards FAO. Upon infection with PVM, we observed an increase in circulating free fatty acids and KB as well as transcriptional reprogramming of the heart supporting FAO (**Figure S1c,d and Figure S3e**). In line with these data and those derived from the LCMV Cl13 infection model, etomoxir treatment of mice infected with PVM led to the predefined humane end points within five hours (**Figure S7e c**). This was accompanied by systemic hypoglycaemia and increased concentrations of markers of tissue damage (**Figure S7f**), corroborating a profound bioenergetic dependency on FAO in independent infection models.

Having observed a strong FAO dependence, we aimed to unveil the systemic and organ-specific bioenergetic changes induced by blockade with etomoxir. To determine effects on systemic FAO, infected mice treated with etomoxir or its vehicle received a dose of ^13^C_16_- palmitate and were analysed for labelled ^13^CO_2_ in metabolic cages. Animals pretreated with vehicle showed an enrichment in labelled ^13^CO_2_, corroborating FAO engagement in the infected host (**Fig. 6e, f**). In contrast, mice pretreated with etomoxir showed a reduced enrichment (**Fig. 6e, f**), demonstrating a systemic failure to oxidize the exogenously administered fatty acid palmitate. Notably, the administration of etomoxir resulted in a quick decline in the mobility of infected mice (**Figure S8a**) and their energy expenditure (**Figure S8b**), indicating a systemic hypometabolic state. Together, these results highlight a relationship between systemic blockade of FAO and reduced survival to infection presumably due to inadequate energy supplies.

Our indirect calorimetry data showed a spike in the respiratory exchange ratio (RER) upon etomoxir injection (**Fig. 6g**), which fell in a range indicative of predominant carbohydrate oxidation^30^. To study this at an organ specific level we administered [^18^F]FDG two hours after etomoxir injection and tracked systemic glucose uptake by whole-body PET/CT imaging. Inhibition of FAO promoted a sharp increase in glucose uptake in virtually all organs including the heart (**Fig.6h, i**), and was more exacerbated in the tissues of infected mice. The spleen was the only organ showing decreased glucose uptake after etomoxir injection (**Fig. 6 h, i**). To address whether FAO blockade affected not only the uptake but also the utilization of energy substrates, we administered a dose of ^13^C_6_-glucose to mice pretreated with etomoxir or vehicle and performed metabolite tracing by mass spectrometry. The spleens of etomoxir-treated animals showed no perturbations in glycolysis, however the contribution of glucose to TCA metabolites was increased in infected animals that received etomoxir (**Fig. 6j**). In the hearts of uninfected mice treated with etomoxir we observed a moderate increase in glucose-derived metabolites of the TCA and glycolysis pathways (**Fig. 6k**), whereas in infected mice, etomoxir treatment dramatically increased the utilization of glucose for synthesis of energy intermediates (**Fig. 6k**). These data indicates that FAO blockade prompts systemic glucose utilization with a high impact on the heart during viral infection.

Heart functionality is highly dependent on a constant supply of energy substrates for ATP synthesis supported mostly by FAO^31^. Two hours after etomoxir administration, i.e. before the animals reached the humane end point, we quantified a significant drop in heart ATP content (**Fig. 6l**), as well as in the energy charge (**Fig. 6m**) in infected animals, but not in uninfected animals. Thus, despite increased oxidation of glucose upon etomoxir administration, this was insufficient to maintain energy production, further corroborating the strong dependence of the heart on fatty acids.

Next, we investigated whether etomoxir injection caused heart damage ^32^. Two hours upon drug injection in mice infected with LCMV Cl13, we did not find increased cell death in heart sections (**Figure S8c**) nor significant changes in parameters of heart contractility such as heart rate (HR), left ventricle ejection fraction (EF) and fraction shortening (FS) (**Figure S8d**). Yet, we observed a trend towards a decline in left ventricular systolic pressure (LVSP), indicating impaired function (**Fig. 3n**). This was accompanied by increased left ventricular end diastolic pressure (LVEDP) observed in LCMV Cl13 infected mice upon etomoxir treatment (**Fig. 6n**), suggesting the progression of LV diastolic dysfunction. These data support the notion that the heart increases its reliance on fatty acids to maintain energy production and heart fitness during times of infection-induced energy substrate redistribution.

### Reduced survival upon FAO blockade is prevented in mice lacking CD8^+^ T cells or lacking GLUT1 in the T cell compartment

We next addressed the question whether CD8^+^ T cell depletion is sufficient to prevent the lethality of FAO blockade during viral infection. Animals receiving anti-CD8 antibody or matched isotype control were infected with LCMV Cl13 and, eight days later, both groups received a single dose of etomoxir. Consistent with the role of CD8^+^ T cells in promoting energy adaptations viral infection, mice lacking CD8^+^ T cells were found to be protected against lethal FAO blockade (**Fig. 7a**) and did not develop severe hypoglycaemia or altered blood levels of KB (**Fig. 7b**). In line with this, *Rag2*^-/-^ mice, or mice depleted of CD8^+^ T cells by antibody, infected with PVM were partially protected against the lethality of FAO blockade (**Figure S9a,b**). Of note, these mice were protected against severe hypoglycaemia but not against hypoketonemia (**Figure S9c**). Together, this data establishes the adaptive immune system as an essential regulator of bioenergetic adaptations during viral infection.

**Fig. 7:**
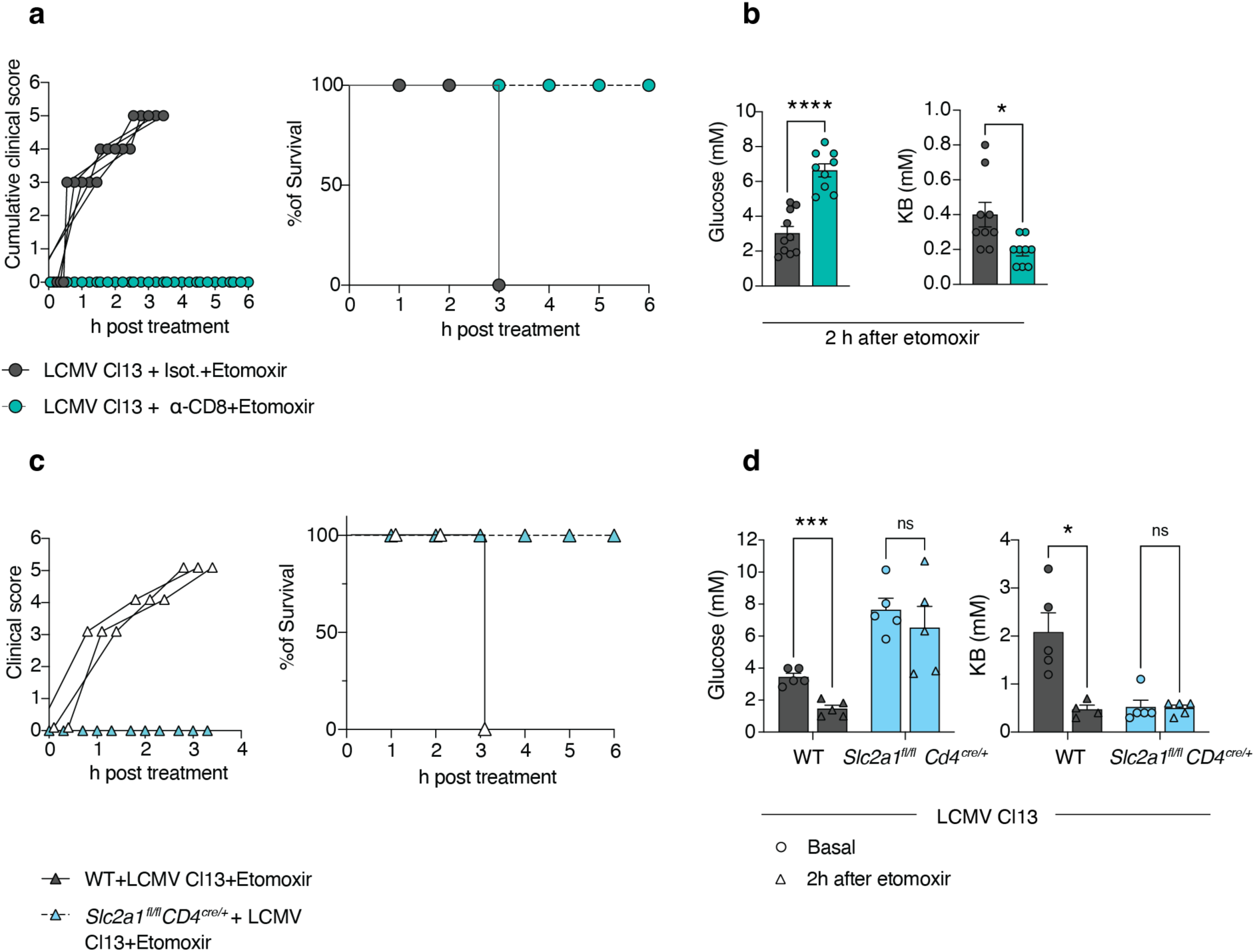
T cells and their glucose uptake are associated with lethal susceptibility to FAO blockade. **a**, Mice were treated with a blocking antibody against CD8 or an isotype control before LCMV Cl13 infection. Mice received one injection of etomoxir eight days after the infection; sickness symptoms were recorded every hour. N= 5 mice per group, the data is representative of two independent experiments. **b**, Glucose and ketone bodies were measured two hours after drug injection. N=9 mice per group, data pooled from two independent experiments. **c**, WT or *Slac2a1^fl/fl^ CD4 ^cre/+^* mice were infected with LCMV Cl13, eight days later mice received an injection of etomoxir and sickness symptoms were recorded every hour. **d**, Blood glucose and KB were measured before and two hours after the injection of etomoxir. N=4-5 mice per group, data pooled from two independent experiments. Unpaired T test. The data represent the mean ± s.e.m. \**p* <0.05, \*\*\**p* <0.0005, \*\*\*\**p* <0.0001.

To assess whether GLUT1 deficiency in the T cell compartment conferred protection against etomoxir-mediated lethality, we injected a single dose of the drug into *Slc2a1 ^fl/fl^ CD4 ^cre/+^* mice and control groups, either of which were infected with LCMV Cl13 eight days before. *Slc2a1 ^fl/fl^ CD4 ^cre/+^* mice were protected unlike the control animals, which gradually progressed to a pre-defined humane endpoint according to clinical symptoms (**Fig. 7c**). *Slc2a1 ^fl/fl^ CD4 ^cre/+^*mice did not develop severe hypoglycaemia, and KB levels remained unchanged upon etomoxir injection in contrast to wildtype mice (**Fig. 7d**). Altogether, these data provide further evidence that the energetic demands of CD8^+^ T cells during infection trigger important systemic metabolic adaptations.

### Spleen and heart bioenergetic reprogramming are observed upon T cell activation in tumour-bearing mice and patients

Finally, we wondered if perturbations in energy substrate utilization can also be observed in other contexts of T cell activation ^24–26^. For this, we retrospectively analysed whole body [^18^F]FDG images of a prospective clinical study of patients with unresectable melanoma. Measurements were conducted before the start of immune check point blockade (ICB) therapy, two weeks and three months thereafter. [^18^F]FDG uptake increased in the spleen two weeks after initiation of ICB compared to the baseline^33^ (**Figure S10a**), and coincided with reduced glucose uptake in the myocardium (**Figure S10b**). Of note, uptake values in both organs returned to basal levels after three months (**Figure S10a,b**). Similar observations were obtained in a retrospective analysis of mice with endogenous insular cell carcinoma treated with ICB in combination with adoptively transferred tumor antigen-specific Th1 cells^34^. There, animals increased uptake of [^18^F]FDG in the spleen while [^18^F]FDG uptake was reduced in the heart seven days after receiving therapy (**Figure S10c,d**). Together, we conclude from these data that the uptake of glucose in spleen and heart may be differentially regulated in the context of T cell activation, irrespective of whether immune activation is triggered by infection.

## Discussion

We discovered an unexpected role for T cell responses regulating energetic metabolism in lymphoid organs and the heart during viral infection. The activation of the immune system was associated with transiently reduced glucose availability, increased fatty acid utilization and corresponding systemic bioenergetic adaptations, particularly affecting the heart.

The shift of the heart to preferential FAO observed in our study may not only reflect an adaptation to low energy availability but could also constitute a protective mechanism to endure infections. For instance, fasting metabolism is associated with resistance to oxidative stress, inflammation, improved metabolic flexibility and contributes to prolonged lifespan, presumably by promoting cellular processes such as autophagy and the production of KB^35,36^. Further, ketogenesis and oxidation of ketone bodies has been associated with the development of protective immune responses^37,38^. It is tempting to speculate that the enforcement of these bioenergetic adaptations linked to improved survival could partially explain why the sick organism reduces food intake during viral infection^39,40^. We hypothesize that the re-allocation of energy substrates tailored to infection is part of the host mechanisms that ensure effective pathogen control, disease tolerance and survival.

We uncovered a crucial role of CD8^+^ T cell - mediated energy re-allocation across organs during viral infection. This appeared to be independent of classical perforin-mediated cytotoxicity and the number of tissue-infiltrating CD8^+^ T cells. Instead, the analysis of mice with T cells harbouring a genetic deficiency for the glucose transporter *Glut1* indicates that glucose utilization by T cells impacts systemic metabolism. The lack of *Glut1* expression in T cells had a moderate impact on T cell activation and a more pronounced effect on energy substrate abundance. Of note, serum levels of IFNγ, TNFα and granzyme B were unaffected in these mice, supporting the notion that GLUT1-mediated uptake of glucose by CD8^+^ T cells, and not these inflammatory mediators, is crucial in impacting the systemic bioenergetic adaptations to viral infection. The underlying mechanisms of T cells orchestrating systemic energy reprogramming could be far more complex than substrate competition and may involve a combination of inflammatory mediators and/or cell-to-cell interactions not explored in this work.

We observed altered glucose uptake in spleen and heart in mouse models of infection as well as in a cancer mouse model and in melanoma patients undergoing ICB. This indicates that adaptations in energy metabolism during immune activation, involving the lymphoid organs and the heart, extend to different underlying stimuli and species. It stands to reason that the relative importance of individual cellular and molecular mediators of bioenergetic adaptations driven by immune activation may vary based on the etiological context, the inflammatory milieu and its progression over time.

The observed reliance on fatty acid utilization during infection was associated with death upon acute FAO blockade, which was preceded by left ventricular failure and metabolic decompensation. This phenotype is reminiscent of patients suffering from inborn FAO disorders, whose fitness and cardiac function can deteriorate rapidly upon acute illness, infection, prolonged fasting and other metabolic stressors ^41^. A strong reliance of the heart on fatty acids has also been implicated in heart failure associated with obesity and type 2 diabetes^3^. These diseases are characterized by an impaired ability to switch energy substrates in organs such as the heart and lead to higher morbidity and mortality during viral infections^4–6,42^. Generally, changes in cardiac energy metabolism are observed in different types of heart failure and may come with complex and different metabolic outcomes^43^. Whether the inability of a particular human organ to adapt its energy demands during infection contributes to these disease outcomes remains unknown. Future research is required to establish whether and how bionergetic adaptations depending on T cells or other immune cells may be causally involved. The host defends itself against infections by deploying pathogen-specific immune responses. There may also be pathogen-specific bioenergetic adaptations necessary for clearance and recovery. We observed similar host changes in the LCMV Cl13 and PVM infection models, affecting both organism-wide and heart-specific glucose and fatty acid metabolism, primarily driven by T cell activation. Yet, LCMV Armstrong and influenza virus infections showed no significant changes in systemic glucose and ketone body levels despite the known activation of T cell responses. This discrepancy is likely to be explained by additional pathogen-specific factors and the ensuing immunometabolic host response. Importantly, LCMV Cl13 infection causes the wasting syndrome cachexia while LCMV Armstrong and influenza infections do not^24,44^. This exemplifies the exquisite specificity of bioenergetic adaptations tailored to pathogens.

This study tackled the fundamental question of how the organism copes with the increasing energy demands of the immune system during viral infection. We show that bioenergetic reprogramming of the heart safeguards its vital function while enabling the immune system to mount an antiviral response. However, this made the heart and the whole organism vulnerable to FAO perturbations. Studying such bioenergetic shifts across organ networks in different contexts of disease will deepen our understanding of molecular pathophysiology on the organismal level and open new avenues for immunometabolic therapies against infectious and inflammatory diseases.

## Non-author contributions

We thank the Biomedical Sequencing Facility from CeMM for sequencing and Gerd Kager for the assistance with the high energy phosphate measurements (Medical University of Graz). We wish to thank Alexander Bartelt, Christoph Binder, Lukas Flatz and Arwand Haschemi for critical discussions and input.

## AUTHOR CONTRIBUTIONS

L.A.H. Conceived the project, designed, and performed experiments, analysed data, interpreted the results and wrote the paper.

A.B. Conceived the project and designed experiments, interpreted the results and wrote the paper.

C.P., S.M.P., X.L. U.U., V.W. and T.W., performed imaging experiments and analysed the data.

A.K., L.W. and C.D. performed and analysed heart functionality experiments.

H.B. designed etomoxir treatment protocol and performed LCMV infection experiments.

1. J. S. and S.K. performed and analysed metabolomics data.

H.C., performed influenza and PVM infection experiments.

F.C. R. and M. S., performed flow cytometry.

J.W and C. V. performed mouse calorimetry experiments.

H.W and S. Ho. Performed in vitro experiments for the *Glut1* KO model.

F.A. analysed bioinformatics data.

S.H. analysed and interpreted the high energy phosphates content of the heart.

E.D.L and C.B. performed PVM infection experiments.

J.L., performed LCMV infections in *Pr1f*1^-/-^KO mice

P.S. performed mouse infection experiments.

F. S. was involved in the retrospective patient study.

B.S. and A.W., were involved in the retrospective mouse tumor study.

A.B., A. H., C.V., performed focus forming assays and qPCRs.

B.D. analysed correlations between PET imaging, viral loads and T cell infiltration.

A.L. and J.W.G. provided help with experiments.

A.O., R.M, A.Q. and J.H. contributed to histological analysis and data interpretation.

M.S., C.C, and T.S., M.V., K. L., M. K., R. Z., S.K., B.N.L., B.P., and M.H., contributed to experimental design and data interpretation.

## METHODS

### Mice

C57BL6/J mice were infected intravenously with 2 x 10^6^ FFU of LCMV Cl13, or LCMV armstrong^20,45^,intratracheally with a sub-lethal dose of 35 PFU of PVM virus strain J3666^22^, or intranasally with a LD50 of influenza A/PR8/34^46^.

For pair feeding experiments, animals were single caged and received a daily amount of food equivalent to the average amount of food consumed by infected animals on a per day basis.

Drugs to modulate glucose (500 mg/kg 2-deoxyglucose, Sigma^47^) and fatty acid oxidation (20 mg/kg etomoxir, Sigma^48^) were administered on the indicated time points by intraperitoneal injection (i.p.). After administration of these drugs, mice were observed, and clinical score was recorded every hour. These included: piloerection, hunched posture, partially closed eyes, laboured breathing, decreased movement and movement only on provocation. Each symptom received a score of one^49^. Animals were immediately euthanized upon reaching the established human end point with a clinical score of five.

For some experiments mice received i.p. 0.5 mg/kg of anti-IFNγ (XMG1, BioXcell) or isotype control (MOPC-21, BioXcell) one day before infection with LCMV Cl13 and every other day for eight days.

For CD8 depletion, mice received on day -2 and -1 prior to infection 0.5 mg/mouse of either anti-CD8 (YTS169.4, InVivoPlus) or a matched isotype control (rat IgG2b, anti-keyhole limpet hemocyanin, InVivoPlus).

Mice received FTY720 (3 mg/kg^50^, i.p. Sigma) starting at three days post infection, every day till day eight. Infection of *Prf1^-/-^* (purchased from Jackson) with LCMV Cl13 was done at the Institute of Immunology, University of Duisburg-Essen, Germany. *Slc2a1^fl/fl^ CD4^cre+/-^* (Jackson labs strain numbers: 031871, 017336) mice were bred at the Institute for Systems Immunology, Würzburg, Germany.

Animals were sex and aged matched and we did not observe significant differences in any of the readouts when employing males or females. Experiments complied with the animal licenses approved by local authorities: GZ-2020-0.406.011, GZ-2024-0.363.541, EC2020-029 and 81-02.04.2019.A143.

### Tracing of ^13^C_6_-glucose and ^13^C_16_ palmitate

Mice received a dose of either ^13^C_6_-glucose (20 mg/mouse^51^ Sigma) or ^13^C_16_-palmitate/BSA (0.3 mg palmitate/mouse^52^, Sigma) i.p., and were euthanized after 10 or 20 min, respectively. Organs were quickly snap frozen and stored at -80 degrees until further analysis. Whole tissue samples were homogenized using a Bertin homogenizer (4 °C, 3 x 30 s at 6500 rpm, 30 s break between each cycle). Then, samples were centrifuged at 10,000 x g during 10 min at 4 °C. Fifty microliters of supernatant were collected, and a Bligh-Dyer extraction was performed to remove lipids and protein. The upper hydrophilic phase was collected and evaporated under a nitrogen flow to dryness. Fifty microliters of water were added to reconstitute the final extract.

A Vanquish UHPLC system (Thermo Scientific) coupled with an Orbitrap Q Exactive (Thermo Scientific) mass spectrometer was used for the LC-MS analysis. Metabolite separation was performed by reversed phase chromatography employing an UPLC Acquity BEH C18, 1.7 M, 2.1 x 100 mm (Waters) analytical column at a column temperature of 40 °C. As mobile phase A, MS-grade water containing 0.1 % formic acid was used. Mobile phase B consisted of MS-grade methanol containing 0.1% formic acid. The flow rate was set to 400 µL/min, and a 3-step gradient of mobile phase B was applied to ensure optimal separation of the analysed metabolites. The mass spectrometer was operated in ESI-positive and -negative mode, capillary voltage 3500 V (positive) and 3000 V (negative), capillary temperature 253 °C, auxiliar gas temperature 406 °C, sheath gas 46 arbitrary units, auxiliary gas 11 arbitrary units and sweep gas 2 arbitrary unit. MS scan mode at 140000 mass resolution was employed for metabolite detection. The scan range was set to 50-450 m/z for both positive and negative ionization modes, the AGC target was set to 1 x 10^6^. Data analysis was performed using TraceFinder software (ThermoFisher Scientific) based on the area of protonated or deprotonated metabolite theoretical masses for molecular ions and isomers with the different heavy carbon atoms incorporated. Quantification of target metabolites was performed with an 8-points calibration curve based on the signal of real neat standards.

### Mice calorimetry

Mice were infected and after eight days treated with etomoxir. Immediately after, mice were individually place in the PhenoMaster (TSE systems) cages. After one hour, mice received 0.3 mg/mouse of ^13^C_6_-palmitate/BSA i.p. Mice were monitored every hour for sickness symptoms. Activity counts, respiratory exchange ratio (RER), oxygen consumption, exhaled ^12^CO_2_ and ^13^CO_2_ were recorded by the metabolic cages for three hours.

### Histology

Whole organs were fixed in 4% paraformaldehyde and paraffin-embedded. 2 µm FFPE organ sections were stained with Hematoxylin (Merck, Darmstadt, Germany) and Eosin G (Carl Roth). Images were photographed using an Olympus BX 53 microscope and adjusted for white-balance.

### Biochemistry

Blood was collected by tail vein nick for measurements of glucose and ketone bodies with the FreeStyle Precision Neo glucometer and test strips (Abbot). Additional blood was collected for serum separation which was kept at -80°C until time of analysis. Determination of serum levels of aspartate aminotransferase (AST), alanine aminotransferase (ALT), blood urea nitrogen (BUN), and creatine kinase muscle brain type (CKMB) were done with a Cobas C311 Analyzer (Roche). Non-esterified fatty acid content in serum was measured with a commercial kit (Fujifilm Wako Chemicals, NEFA-HR(2)).

### qPCR

Whole hearts were homogenized using a TissueLyser II (QIAGEN). RNA was extracted using QIAzol following manufacturer’s instructions (QIAGEN). cDNA synthesis was done using random primers and the First Strand cDNA Synthesis kit (Thermo Fischer Scientific). Real time PCR was performed with the Taqman Fast Universal PCR Mastermix (Thermo Fischer Scientific) and Taqman Gene expression Assays (Thermo Fischer Scientific) for *Slc2a4* (Mm00436615_m1), *Slc27a1* (Mm00449511_m1), *Pdk4* (Mm01166879_m1) *Hmgcs2* (Mm00550050_m1), *Cpt1b* (Mm00487191_g1), *Cd36* (Mm00432403_m1) and *Rplp0* (Mm00725448_s1).

### Quantification of viral loads

For LCMV viral load determination a modified focus forming assay was done using Vero cells (ATCC CCL-81)^53^. LCMV NP RNA load was quantified as described previously^20^.

### Positron emission tomography

Non-invasive in vivo imaging was acquired on a Siemens Inveon preclinical µPET/SPECT/CT system (Siemens Medical Solutions, USA). Mice were anaesthetized using 1.5-2 % isoflurane with oxygen (1.5-2 l/min) and warmed (37°C) in the positioning bed. After a µCT scan of 7 min, dynamic PET imaging was acquired for 60 min. Therefore, 11-17 MBq [^18^F]FDG or [^18^F]FTHA were injected intravenously (lateral tail vein) into the immobilized animal. The total volume did not exceed 200 µL. Anaesthesia was maintained during the entire measurement. CT raw data were reconstructed with a Feldkamp algorithm using a Shepp-Logan filter followed by standard mouse beam-hardening correction and noise reduction (matrix size: 1024 × 1024; effective pixel size: 97.56 µm). The dedicated CT image data were calibrated to Hounsfield Units (HU). PET list mode data were sorted into three-dimensional sinograms and reconstructed using an OSEM 3D/OP-MAP scatter corrected reconstruction algorithm (matrix size 256 × 256). The data were normalized and corrected for random, dead time and radioactive decay. A calibration factor was applied to convert the activity information into absolute concentration units. Multimodal (µPET/CT) rigid-body image registration and biomedical image quantification was performed using the image analysis software PMOD 3.8 (PMOD Technologies, Switzerland) and Inveon Research Workplace (IRW; Siemens Medical Solutions, USA). Volumes of interest (VOIs) were outlined on multiple planes of the CT and PET images. Time-activity-curves (TACs) were calculated, normalized to injected dose and animal weight and expressed as standardized uptake values (SUV; g/mL) to facilitate the comparison.

### Transthoracic echocardiography

Left ventricular systolic and diastolic functions was measured non-invasively with conventional echocardiography (Vevo 3100 Imaging System with a 40-MHz linear probe (Visualsonics) as described previously^54^. The animals were anesthetized by 1%–1.5% isoflurane and the ECG was monitored via limb electrodes. Left ventricular ejection fraction (LVEF), end-diastolic and end-systolic diameters were evaluated at a mid-papillary short axis view. For each parameter, a mean of three cardiac cycles in each view was used.

### Assessment of left ventricular hemodynamic function *in vivo*

Left ventricular (LV) hemodynamic parameters were invasively measured as described previously^55^. Mice were anaesthetized by intraperitoneal injection of a mixture of Xylazin (4 mg/kg; Bayer, Germany) and Ketamin (100 mg/kg; Dr E. Gräub AG, Switzerland), intubated and ventilated. The chest was opened and a microtip catheter (SPR-1000, Millar, Millar Instruments, Houston, USA) was gently inserted into the LV chamber. Hemodynamic parameters such as left ventricular systolic (LVSP), left ventricular end-diastolic pressure (LVEDP), and heart rate (HR) were continuously recorded on Labchart (v7.3.2, Powerlab System (8/30)), both AD Instruments, Spechbach, Germany).

### Measurement of high phosphates in the left ventricle

Mice were anaesthetized by intraperitoneal injection of a mixture of Xylazin (4 mg/kg; Bayer, Germany) and Ketamin (100 mg/kg; Dr E. Gräub AG, Switzerland), The chest was opened, and the heart was freeze-clamped (liquid nitrogen). The obtained hearts were either immediately subjected to the work-up procedure or stored in liquid nitrogen prior to determination of high energy phosphates by HPLC as previously described^56,57^. The pellets of the acid extract of the hearts were dissolved in 1 mL of 0.1mol/L sodium hydroxide and further diluted 1:10 with physiological saline for protein determination (BCA Protein Assay, Pierce; Pierce Biotechnology, Rockford, IL, USA). Energy charge was calculated using the formula: (ATP + 0.5ADP) / (AMP + ADP + ATP).

### Acylcarnitines profiling

Liver and serum samples of uninfected and LCMV Cl13 infected mice were collected and snap frozen in liquid nitrogen until analysis. Targeted metabolomics was performed using the AbsoluteIDQ p180 kit (Biocrates Life Sciences AG, Innsbruck, Austria). The samples were analyzed on an AB SCIEX QTrap 4000 mass spectrometer using an Agilent 1200 RR HPLC system, which were operated with Analyst 1.6.2 (AB SCIEX). The chromatographic column was obtained from Biocrates. The serum samples and additional blanks, calibration standards and quality controls were prepared according to the user manual. The experiments were validated with the supplied software (MetIDQ, Version 5-4-8-DB100-Boron-2607, Biocrates Life Sciences, Innsbruck, Austria).

### Seahorse extracellular flux analysis

Mitochondrial respiration and aerobic glycolysis of CD8^+^ T cells was measured as their oxygen consumption rate (OCR) and glycolytic proton efflux rate (PER), respectively, using an oxygen-controlled XFe96 extracellular flux analyzer (Seahorse Bioscience). CD8^+^ T cells were stimulated on 12-well delta-surface plates (Nunc). The plates were pre-coated with 12 μg/ml polyclonal anti-hamster IgG (MP Biomedicals) for 2 h, washed once with PBS and 2x10^6^ T cells were activated with plate-bound 0.5 μg/ml of anti-CD3 (clone 145-2C1) and 1 μg/ml anti-CD28 (clone 37.51, both Bio X Cell) for 24 and 48h. XFe96 cell culture microplates (Agilent) were coated with 22 μg/ml Cell-Tak (Corning) and 2x10^5^ T cells per well were attached in 4-8 replicates in Seahorse XF RPMI medium (Agilent) supplemented with 2 mM L-glutamine (Gibco), 1 mM sodium pyruvate (Sigma) and 10 mM D-glucose (Sigma). After 1h incubation in a CO2-free incubator at 37°C, glycolytic and mitochondrial stress tests were performed according to the manufacture’s recommendation. In brief, for assessing glycolysis, basal extracellular acidification rate (ECAR) was measured followed by addition of 0.5 μM rotenone (AdipoGen) and 0.5 μM antimycin A (Sigma) to inhibit mitochondrial complex 1 and 3, respectively. At the end of the measurement, 50 mM 2-DG (Sigma) was added to block glycolysis. To analyze mitochondrial respiration, basal oxygen consumption was measured followed by the addition of 2 μM oligomycin (Cayman Chemicals), an ATP synthase inhibitor, 1 μM of the protonophore carbonyl cyanide-*4*-(trifluoromethoxy)-phenylhydrazone (FCCP, Cayman Chemical) and 0.5 μM rotenone (AdipoGen) together with 0.5 μM antimycin A (Sigma). The basal oxygen consumption was calculated by subtracting the OCR after rotenone and antimycin A treatment from the OCR before oligomycin treatment. The maximal OCR was calculated by subtracting the OCR after rotenone and antimycin A treatment from the OCR measured after addition of FCCP.

### Tissue Preparation and Immune Cell Isolation

Transcardiac perfusion of mice was done with ∼20mL PBS. Splenocytes were isolated by straining cells through 70 µm strainers. Cells were subsequently incubated with 1x RBC lysis buffer (eBioscience, #00-4333-57) for 5 min at room temperature. Cells were washed then subjected for flow cytometry staining.

Liver immune cells were isolated by mashing the tissue through 70 µm strainers. Cells were then resuspended in 5 mL of a 42% Percoll solution and centrifuged for 20 min at 850 g with break off. Cell pellet was recovered and then incubated with 500 µL 1x RBC lysis buffer for 3min at room temperature. Cells were washed and then subjected for flow cytometry staining.

Kidney immune cells were isolated by mincing the tissue in 3mL digestion buffer containing PBS, 2% FBS and 1mg/mL Collagenase B (Roche, # 11088807001). Digestion solution was then incubated for 45 min at 37°C under agitation. Tissue pieces were then disrupted by pipetting them using 1mL wide-bore tips several times. Cell suspension was strained through 70 µm strainers. Cells were then resuspended in 5 mL of a 42% Percoll solution and centrifuged for 20 min at 850 g with break off. Cell pellet was recovered and then incubated with 500 µL 1x RBC lysis buffer for 3 min at room temperature. Cells were washed and then subjected for flow cytometry staining.

One hemisphere was used for immune cell isolation from the brain. Brain was minced and digested in 2mL digestion buffer containing RPMI, 10% FBS and 0.2 mg/mL Liberase TL (Roche, #05401020001) for 30 min at 37°C under agitation. Cell suspension was gently disrupted by pipetting up and down using 1 mL widebore tips. Cells were subsequently resuspended in 5 mL of 42% Percoll solution and centrifuged for 20 min at 850 g with break off. Cell pellets were washed and then subjected for flow cytometry staining.

Adipose tissue digestion was performed as previously described^58^. In brief, adipose tissues were collected in DMEM containing 1% fatty-acid free BSA. Tissues were subsequently minced and digested in the DMEM containing 1% fatty-acid free BSA, 0.2 mg/mL Liberase LT and 10 µg/mL DnaseI (Roche, #11284932001) for 30 min at 37°C under agitation. Tissues were disrupted by pipetting using 1mL widebore tips and strained through 70 µm strainer. Cell pellets were washed and then subjected for flow cytometry staining.

Heart and quadriceps were isolated and collected in cold PBS. Tissues were minced and digested in RPMI, 10% FBS and 725 U/mL Collagenase type II (GIBCO, #17101-015) for 25 min at 37°C under agitation. Tissues were gently disrupted by resuspending using 5 mL serological pipettes and then strained through 70 µm strainer. Cell pellets were washed and then subjected for flow cytometry staining.

### [^3^H]-2-DG uptake of activated CD8^+^ T cells

Glucose uptake was directly assessed using tritiated 2-deoxy-glucose ([3H]-2-DG) from Perkin Elmer. CD8^+^ T cells were activated with 0.5 mg/ml anti-CD3 (clone 145-2C1) and 1 mg/ml anti-CD28 (clone 37.51, both from Bio X Cell) for 24. Following activation, 5x10⁵ T cells were incubated in 200 µl of glucose-free medium (supplemented with 10% FBS) containing 1 µCi/ml [3H]-2-DG in 96-well plates. Incubation times were 30 minutes, 1 hour, 2 hours, or 4 hours. After incubation, T cells were washed twice with ice-cold PBS, resuspended in 100 µl of H₂O, and 40 µl of the lysate was mixed with 160 µl of scintillation cocktail (Roth). Intracellular [3H] counts per minute (cpm) were measured using a MicroBeta2 microplate scintillation counter (Perkin Elmer) and normalized to the cell count.

### Flow cytometry

Cells were stained with Fixable Viability Dye eFluor 780 (eBioscience) for 15 min in PBS at RT together with an anti-FcgRII/FcgRIII antibody (clone 2.4G2; Bio X Cell) to prevent unspecific binding. After washing, surface antigens were stained using fluorophore-conjugated antibodies in PBS containing 0.5% BSA for 20 min at RT in the dark. For detection of GLUT1 protein, stimulated CD8^+^ T cells were fixed with IC-fixation buffer (eBioscience) and GLUT1 was stained intracellularly in 1x permeabilization buffer (eBioscience) with a rabbit polyclonal anti-mouse GLUT1 antibody (Abcam, EPR3915) diluted 1:100 in permeabilization buffer o/n (eBioscience). After washing, the primary antibody was detected using an anti-rabbit IgG secondary antibody conjugated to Alexa Fluor 546 (Invitrogen) diluted 1:1000 in permeabilization buffer for 40 min. All sample acquisition was performed with BD Celesta (BD Biosciences) flow cytometers and further analysed with the FlowJo software (BD Biosciences).

### Retrospective data analysis of a patient study

We evaluated a subcohort of patients recruited in a prospective study approved by the local ethics committee (code: 251/2012BO1), the Federal Agency for Radiation Protection (code: Z5-22463/2-2012-023), registered on clinicaltrial.gov (NCT 03132090) and the German clinical trials register (DRKS00013925). Results of this study have been published previously and technical details can be found in^33,59–61^. Briefly, the study includes adult patients with unresectable metastasized melanoma, scheduled for immunotherapy. Whole-body ^18^F-FDG-PET/MRI scans were performed just before planned treatment initiation (t_0_), two weeks (t_1_) and three months (t_2_) after therapy start. Written informed consent was given by all patients. To evaluate changes of the glucose consumption (measured as SULmean) of the heart and spleen, we performed a new analysis of patients included in^33^. Patients with an SULmean of the heart at t_0_ <1 were excluded from further evaluation. Finally, 14 patients met the inclusion criteria (table 1^33^) and the SUVmean of the heart and the spleen were measured at all three time points.

### Retrospective analysis of mouse model of insular cell carcinoma with immunotherapy

The data analysed was previously published and detail can be found in^62^. Briefly, transgenic Rip1-Tag2 mice bearing insular cell carcinomas were transferred into a radiation cylinder for a preparative immune cell depleting low-dose whole body radiation (dose: 2 Gy) 24 h before intraperitoneal (i.p.) weekly transfer of 10^7^ tumor-antigen (Tag2) specific T_h_1 cells. Some mice remained non-radiated. For immune checkpoint blockade (ICB), anti-PD-L1 mAb (clone 10F.9G2, Bio X Cell) and anti-LAG-3 mAb (clone C9B7W, Bio X Cell) were i.p. injected twice/week one and four days after T_h_1 transfer (initially 500 μg, afterwards 200 μg per mouse). Mice were fasted overnight prior to ^18^F-FDG injection after 4 weeks of treatment. Mice were anaesthetized with 1.5% isoflurane in oxygen in a temperature-controlled anaesthesia box. 12.7 ±0.5 MBq of ^18^F-FDG dissolved in 150 µL NaCl solution was injected intravenously (i.v.) via the tail vein. The animals were kept warm under isoflurane anaesthesia to allow ^18^F-FDG to reach equilibrium of accumulation in the myocardium. PET scans were performed after 45-60 min of unconscious ^18^F-FDG uptake on a dedicated small-animal PET scanner (Inveon, Siemens Preclinical Solutions, Knoxville, USA) for 15-20 min maintained by isoflurane narcosis. According to our standard protocol for mouse PET imaging, no attenuation correction was applied. Reconstruction was performed in Inveon Acquisition Workplace 1.5.0.28 with OSEM2D with four iterations. ^18^F-FDG uptake in the left ventricle of the heart was analysed using PCARD (PMOD version 4.206, PMOD Technologies Ltd, CH). For this, volumes of interest (VOIs) were created based on PET images and corrected for radioactive decay. The data is expressed as standard uptake value normalized for body weight (SUV_BW_).

### Measurement of cytokines in serum

IFNγ and TNFα were measured in serum samples with the Legendplex mouse anti-virus response panel (Biologend, 740621) according to the manufacturer’s instructions. Granzyme B was measured by ELISA (Invitrogen, 88-8022) according to the manufacturer’s instructions.

### Statistics

Data is depicted as mean ± s.e.m. Statistical significance was calculated as indicated in the respective figure legends using GraphPad Prism 10.

## Supplemental figures

**Extended Data Fig. 1:**
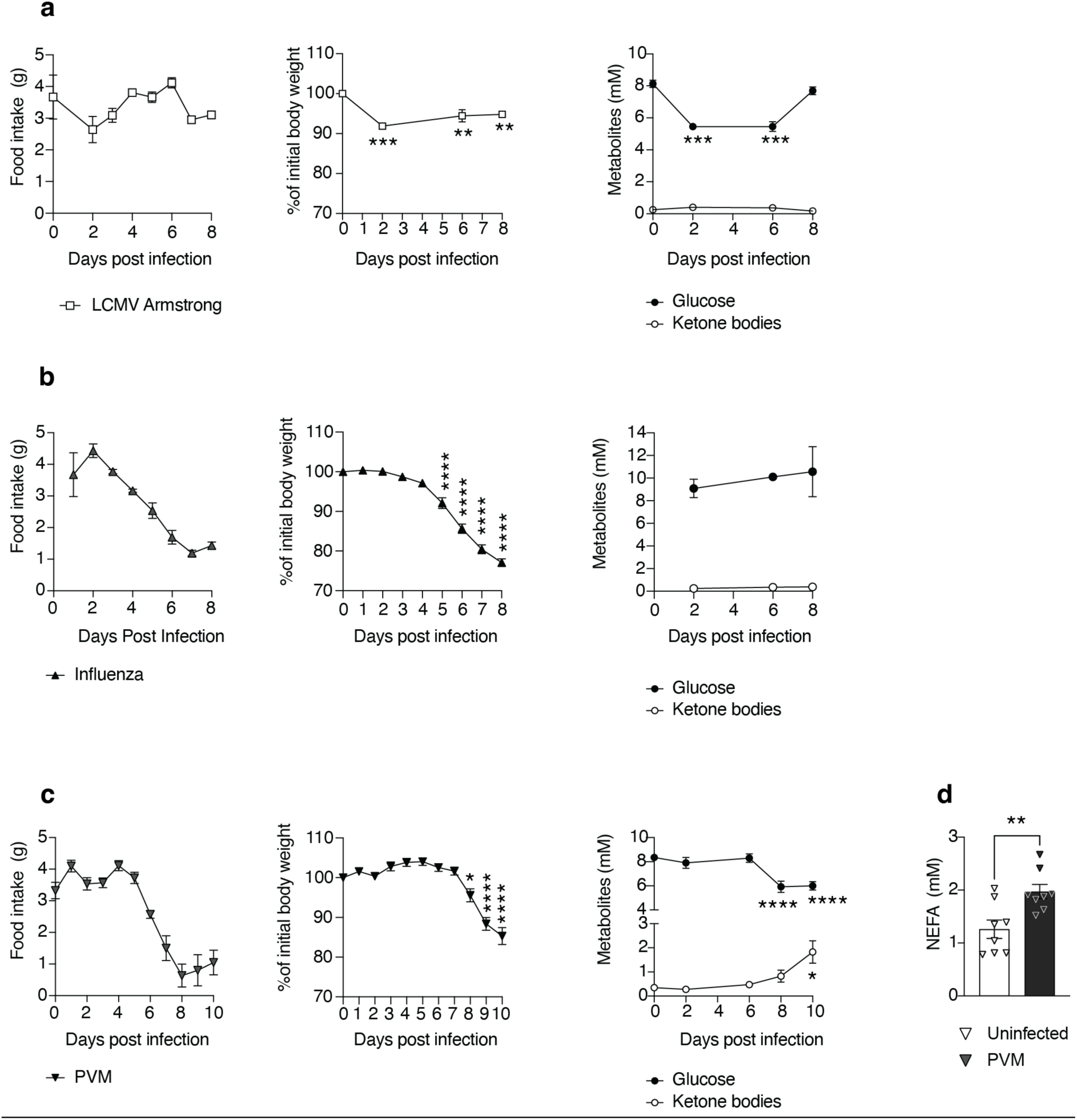
Characterization of metabolic changes in different infection models. Mice were infected with LCMV Armstrong (**a**), Influenza PR8 (**b**) or PVM (**c**). The graphs show food inatek, the percentage of body weight at a given time compared to the weight at the time of infection, and the concentrations of blood glucose and KB. N=4-5 mice per group, the data are representative of at least two independent experiments. Statistical significance against their respective time zero. One-way ANOVA with Bonferroni correction. **d**. Concentrations in serum of non-esterified fatty acids (NEFA) of uninfected or mice infected for 10 days with PVM. N=8 mice per group, data pooled from two independent experiments. Unpaired T test. \**p* <0.05, \*\**p* <0.005, \*\*\**p* <0.0005, \*\*\*\**p* <0.0001.

**Extended Data Fig. 2:**
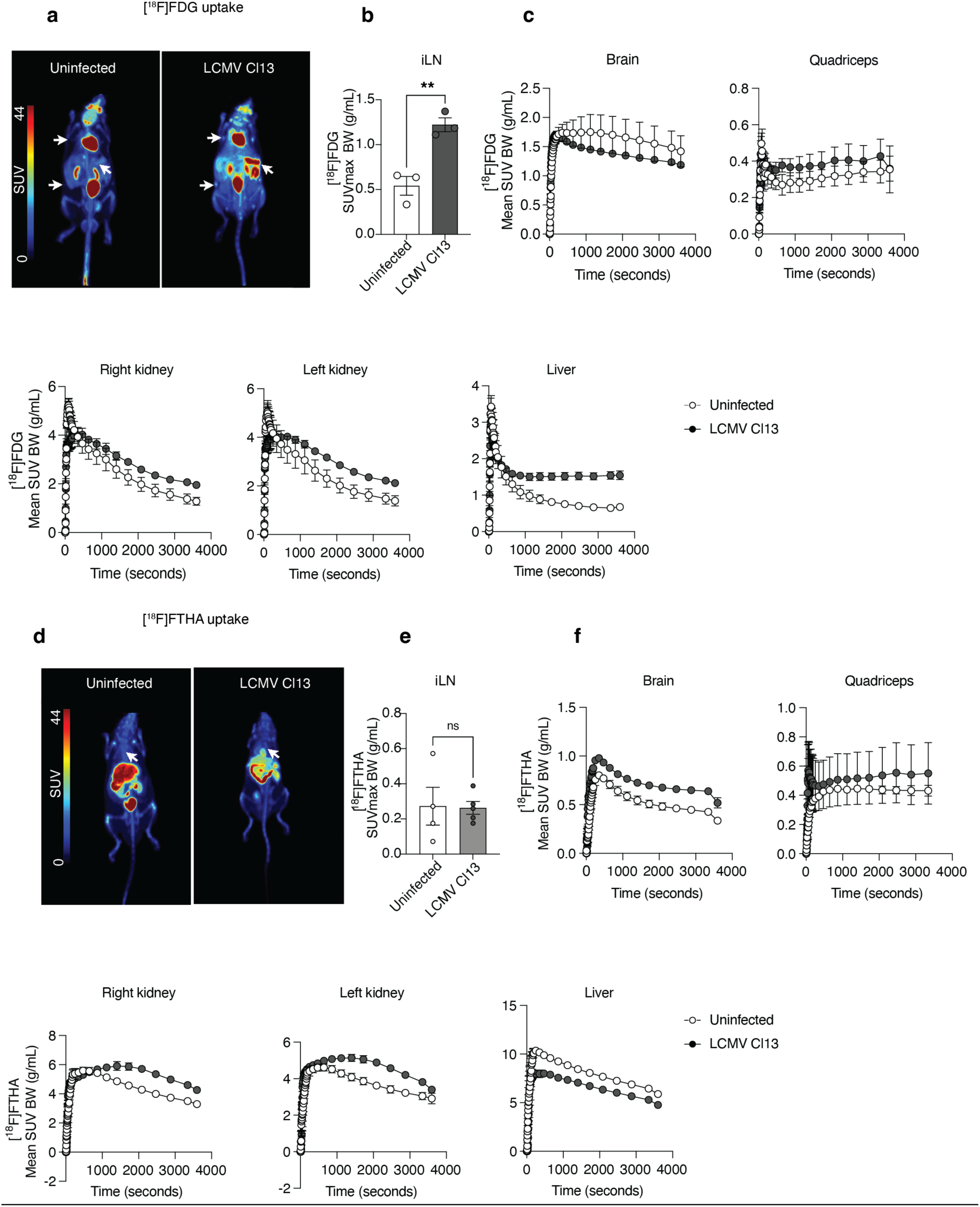
Whole body imaging of glucose and fatty acid uptake. Mice were infected with LCMV Cl13 and after eight days they received an i.v. injection of [^18^F]FDG or [^18^F]FTHA. **a**, Representative image represents tracer uptake in the whole mouse 60 min after injection. **b**, [^18^F]FDG uptake in the right inguinal lymph node after 60 min. **c**, Time activity curves of tracer uptake in different organs. **d**, Representative image of [^18^F]FTHA tracer uptake in the whole mouse 60 min after injection. **e**, Uptake of the tracer in the right inguinal lymph node after 60 min. **f**, Time activity curves of tracer uptake in different organs. In **a**, the arrows on the left point at the axillary and inguinal lymph nodes and on the right side to the spleen. In **d**, the arrows point to the heart. N= 3-4 mice per group, the data are representative of at least two independent experiments. The data show the mean ± s.e.m. Unpaired T test. \*\**p*<0. 005.representative of at least two independent experiments.

**Extended Data Fig. 3:**
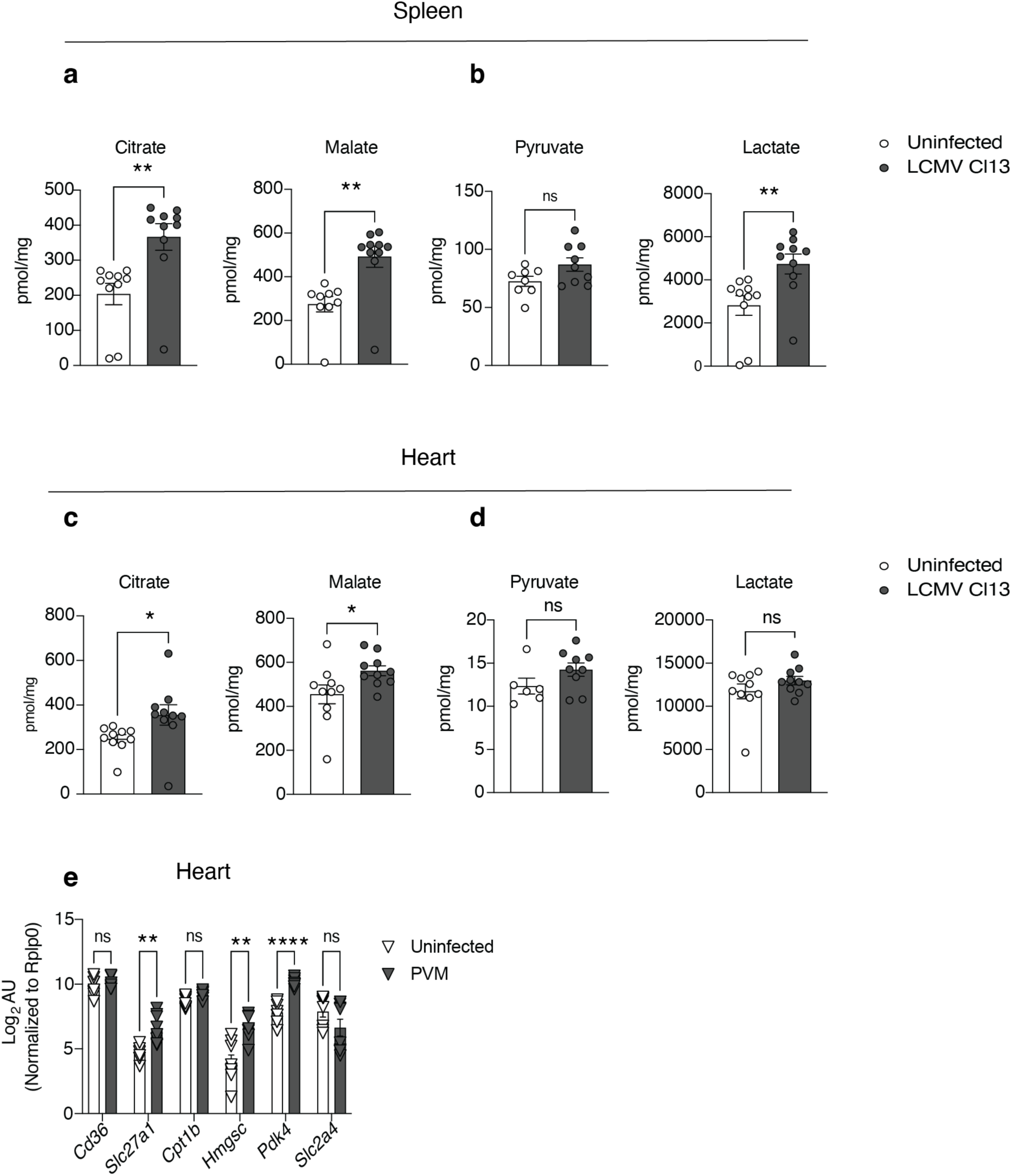
Metabolic changes in the models of LCMV Cl13 and PVM. Organs were recovered from uninfected or mice infected with LCMV Cl13 for eight days for measurement by mass spectrometry of TCA and glycolysis metabolites. **a, b,** shows metabolite concentrations/mg of tissue in the spleen and **c, d,** in the heart. N=6-10 mice per group, data was pooled from two independent experiments. **e**, WT Mice were intratracheally infected with 35 PFU PVM strain J3666. Gene expression analysis by qPCR from hearts were done on day nine after infection. N= 8 mice per group, data pooled from two independent experiments (these data set is also shown in Ext. Data Fig. 4i). The data show the mean ± s.e.m. Unpaired T test. \**p*<0.05, \*\**p*<0.005, \*\*\**p* <0.0005, \*\*\*\**p* <0.0001.

**Extended Data Fig. 4.**
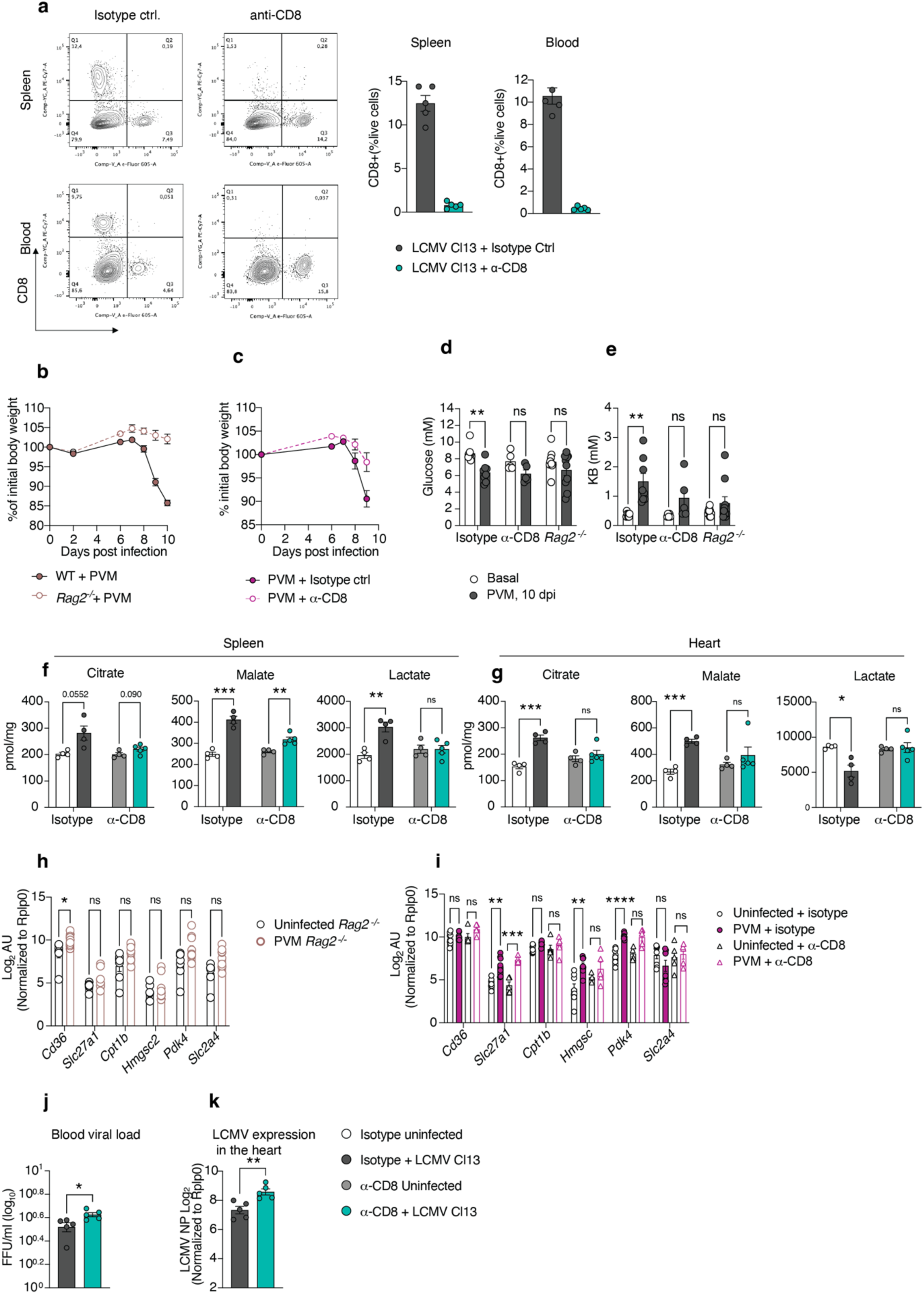
Characterization of infected *Rag2*^-/-^ and CD8 deficient mice. **a**, WT mice received at days -2 and -1, 0.5 mg/mouse of an anti-CD8 antibody or a matched isotype control, thereafter they were infected with LCMV Cl13. The FACS plots are representative of spleen and blood and show the staining of CD4 and CD8 in live cells. The graphs depict the percentage of CD8^+^ cells in both spleen and blood. N= 4-5 mice per group. The data is representative of at least two independent experiments. **b**, WT and Rag2-/- mice were infected with PVM, the graph shows changes in body weight. N=9-10 mice per group, data pooled from two independent experiments. **c**, WT received an anti-CD8 antibody or isotype control as in (**a**) and infected with PVM. N=7-10 mice per group, the data was pooled from two independent experiments. **d**, **e**, shows changes in blood glucose and KB. **f**, Mice treated as in **a** received one injection of ^13^C_6_-glucose eight days after infection with LCMV Cl13. Ten minutes later organs were recovered for quantification of the total pool of TCA and glycolysis metabolites in the spleen and heart. N=4-5 mice per group. **h, i,** gene expression changes in the heart of mice infected with PVM. **j**, Focus forming assay was done using blood samples collected at eight days post infection with LCMV Cl13 and depicts the number of focus forming units (FFU). **k**, LCMV NP expression was estimated by qPCR (right) of samples obtained from heart homogenates. N= 5 mice per group. Data are representative of two independent experiments. The data show the mean ± s.e.m. Unpaired T test. \**p*<0.05, \*\**p*<0.005, \*\*\**p* <0.0005, \*\*\*\**p* <0.0001.

**Extended Data Fig. 5:**
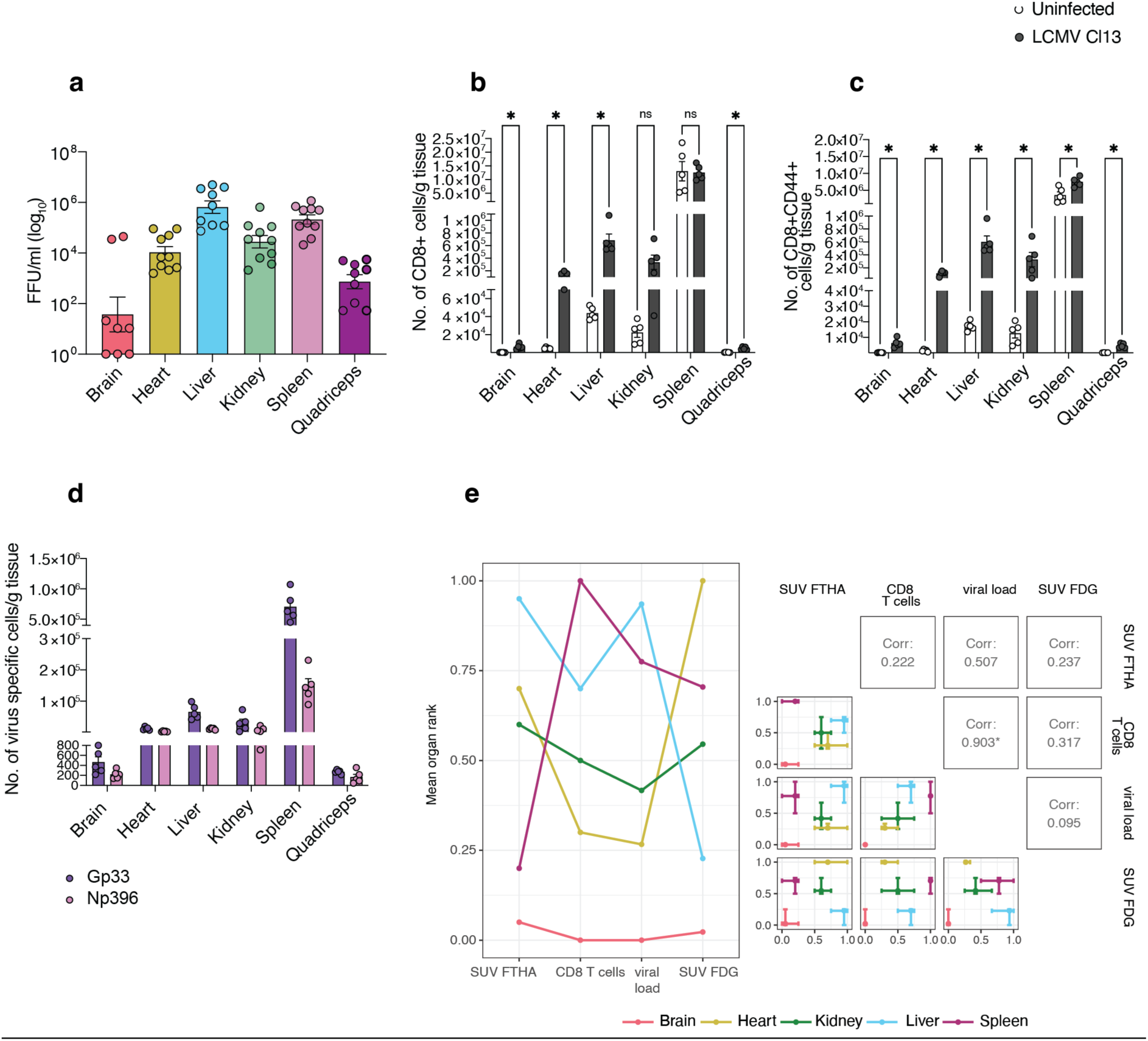
Viral loads and CD8 T cell infiltration across tissues. Mice were infected with LCMV Cl13 and after eight days organs were recovered. **a**, viral loads in different tissues, N=8-10 mice per group, data was pooled from two independent experiments. **b**-**c**, Tissues were weighed, digested and then stained with fluorescent antibodies to quantify **b**, total CD8 T cells, **c**, activated CD44^+^CD8^+^ T cells and **d**, virus specific CD8^+^ T cells. N=5 mice per group, the data is representative of two independent experiments. Unpaired T test. The data show the mean ± s.e.m. **e**, The SUV-FTHA, SUV-FDG (from Fig.1), and T cell measurements were normalised relative to the median of the uninfected values. No normalisation was done for the viral loads. Since measurements were taken in different mice, a consensus rank approach was taken for comparison purposes. For each measurement, organs (brain, heart, kidney, liver and spleen) were first percentage ranked within each mouse, and then the consensus rank for each organ across mice was calculated using the mean. Lower consensus ranks imply lower values measured for that organ. The association of the consensus rankings between pairs of measurements were compared using Pearson’s correlation. The pairwise correlation plot shows the consensus rank (point) alongside the minimum and maximum ranking for each organ (error bars).

**Extended Data Fig. 6:**
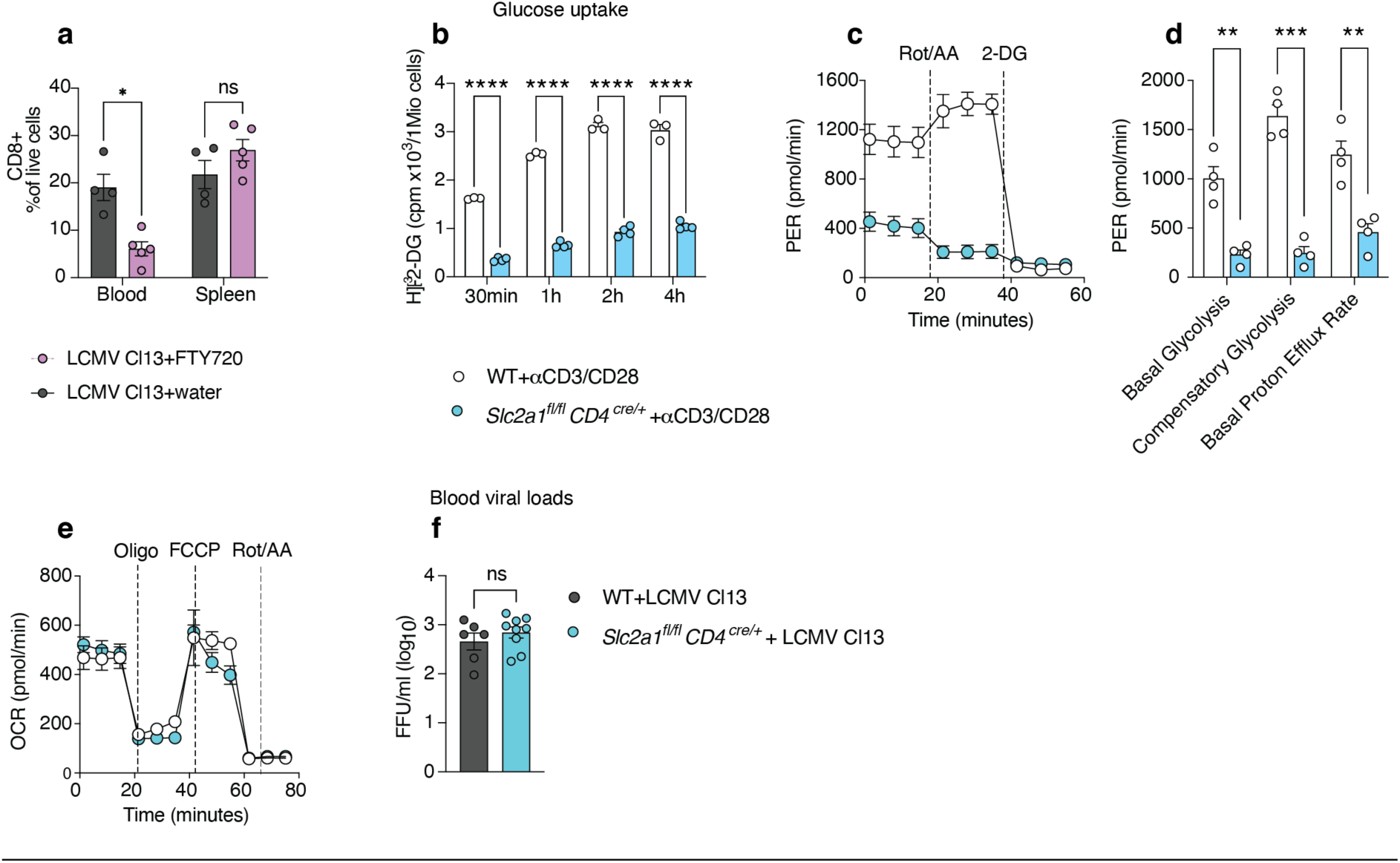
characterization of T cell specific models. **a**, WT mice were infected with LCMV Cl13. After four days they received an injection of FTY720 every day till day eight. The graph shows CD8^+^ T cells in blood and spleen. **b-e,** CD8 T cells were obtained from the spleens of WT and *Slc2a ^fl/fl^ CD4^cre/+^* mice. The cells were activated by plate-bound a-CD3/CD28 stimulation for 24 h followed by measurements of **b**, [^3^H]-2-DG uptake or (**c-e**) extracellular flux analysis. **c, d,** determination of glycolysis capacity and **e**, mitochondrial oxidative phosphorylation. N=4 mice per group. Data are representative of two independent experiments. **f**, WT and *Slc2a ^fl/fl^ CD4^cre/+^* mice were infected with LCMV Cl13, eight days viral loads were measured in blood, N=6-9 mice per group, data was pooled from two independent experiments. The data show the mean ± s.e.m. Unpaired T test. \**p*<0.05, \*\**p*<0.005, \*\*\**p*<0.0005, \*\*\*\**p*<0.0001.

**Extended Data Fig. 7:**
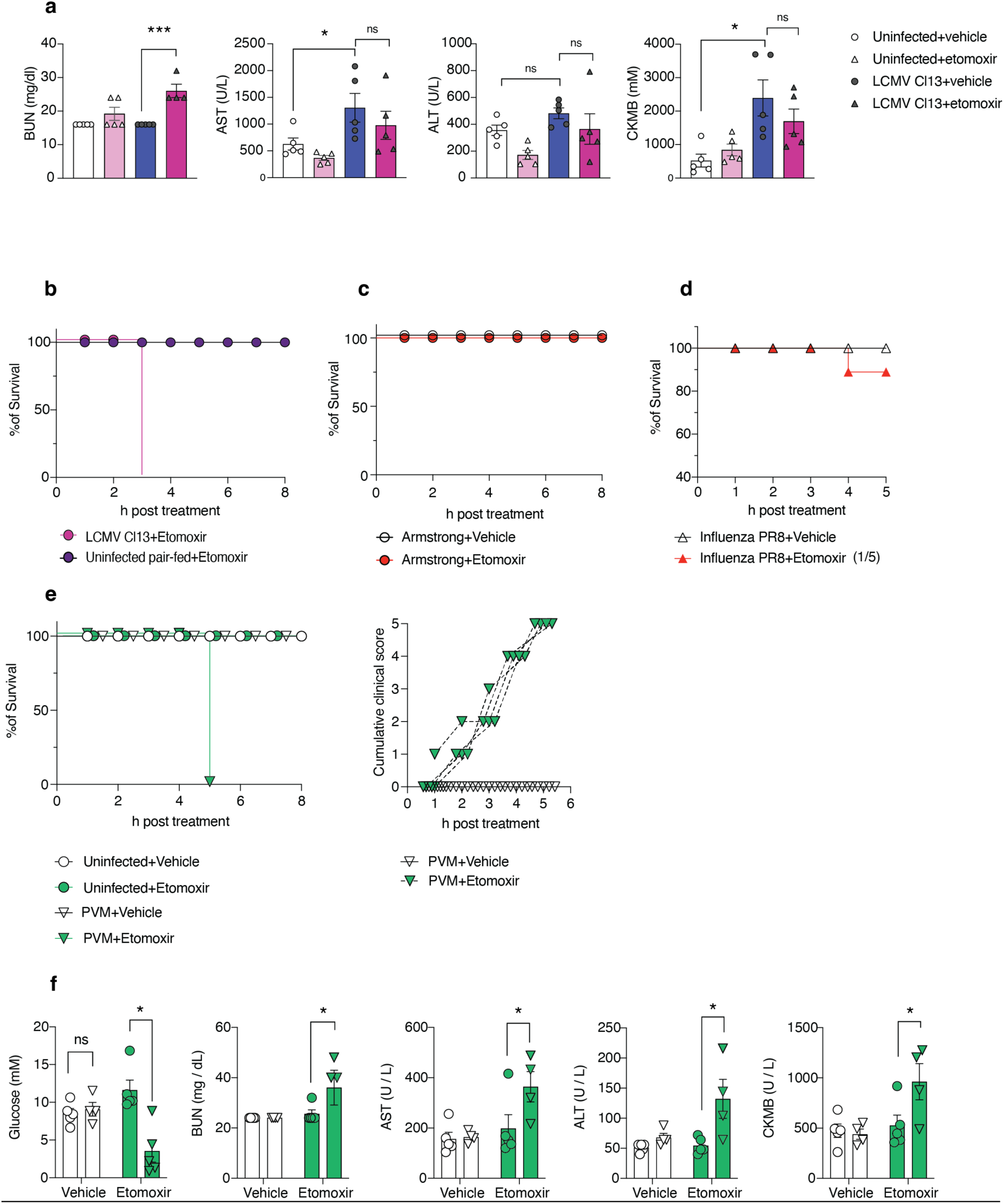
Characterization of mice upon pharmacological FAO blockade. Mice were infected with LCMV Cl13. After eight days they received a single dose of 20 mg/kg of etomoxir or vehicle. **a**, Four hours after drug treatment, serum was collected to measure concentrations of markers of tissue damage. BUN: blood urea nitrogen, ALT: alanine aminotransferase and AST: aspartate aminotransferase, CKMB: creatine kinase isoenzyme MB. N=4-5 mice per group. Representative data of two independent experiments. **b**, pair fed mice (receiving the average daily amount of food consumed by LCMV Cl13 infected mice), or mice infected with **c**, LCMV Armstrong or **d**, influenza PR8 received a single dose of 20 mg/kg of etomoxir or vehicle, eight days post infection. N=5 mice per group, the data is representative of two independent experiments. **e**, WT Mice were intratracheally infected with 35 PFU PVM strain J3666. Survival after a single injection of 20 mg/kg of etomoxir or vehicle at day nine post infection. In the graph, 0% indicates mice reached the humane end point and were euthanized. N= 5 mice per group. The graph on the right shows the cumulative clinical score of infected mice challenged with etomoxir or vehicle. Data is representative of at least two independent experiments. **f**, Upon sacrificing, serum samples were obtained to measure concentrations of glucose and markers of tissue damage. N= 5 mice per group. The data are representative of two independent experiments. The data show the mean ± s.e.m. Unpaired T test, \**p*<0.05, \*\*\**p*<0.005.

**Extended Data Fig. 8:**
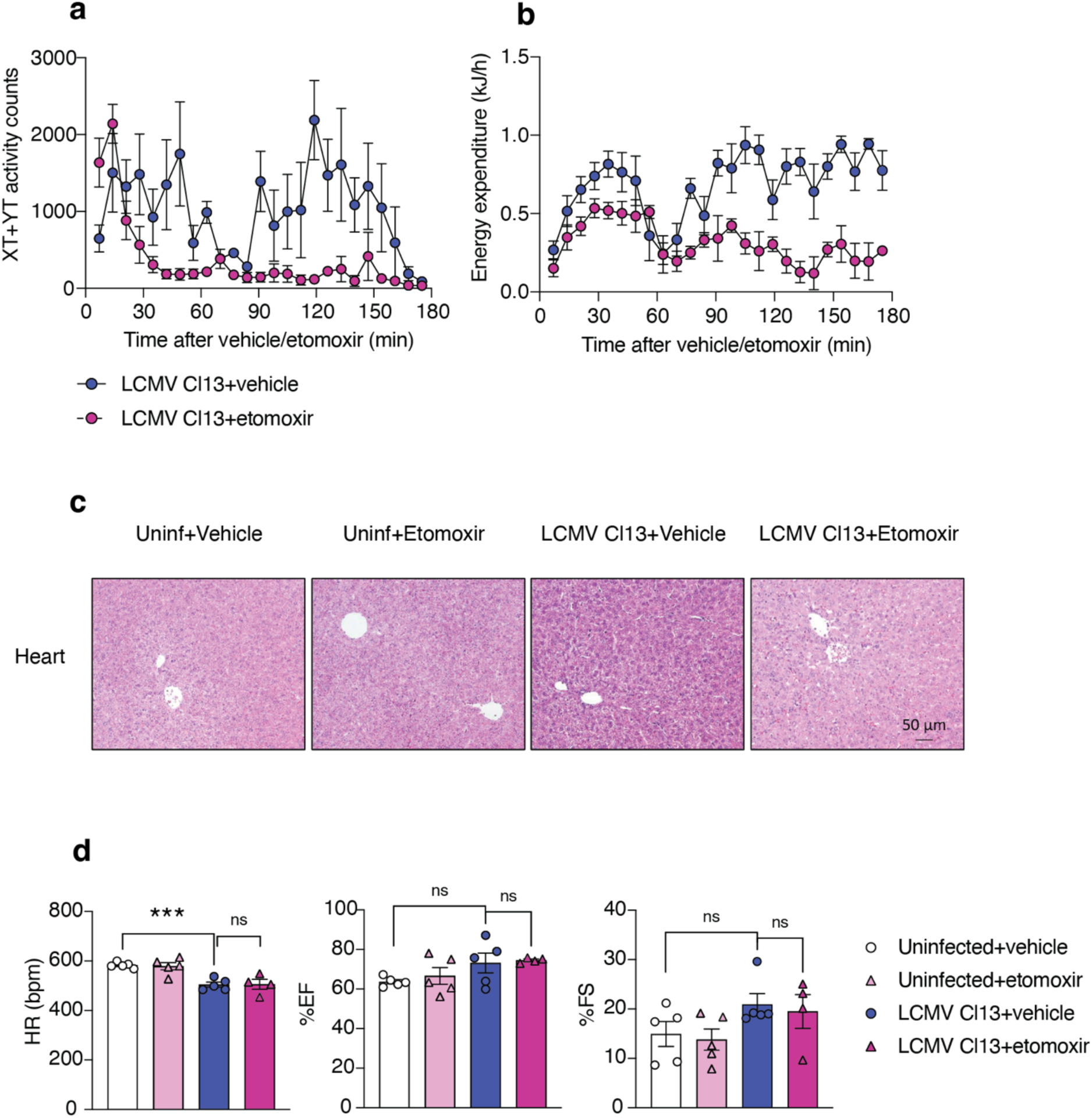
Characterization of pathology upon etomoxir injection in the model of LCMV Cl13. **a**, **b**, Mice received a dose of etomoxir eight days after infection with LCMV Cl13. Immediately after animals were placed in metabolic cages. The graphs show activity counts (**a**) and energy expenditure (**b**) before animals reached the humane end point. N=6 mice per group, data was pooled from two independent experiments. **c,** Hearts were collected two hours after drug treatment of mice infected with LCMV Cl13. H&E staining and histologic analysis were performed. Representative images are showed. N=4-5 mice per group. **d**, Heart functionality of uninfected and infected mice (eight days post-infection) was assessed by echocardiography two hours after vehicle or etomoxir administration. HR; heart rate, EF: ejection fraction, FS: fraction shortening. N= 4-5 mice per group. **e**, concentration of lactate in serum, **f**, ratio of NADH/NAD+ in heart and **g**, concentration of non-esterified fatty acids (NEFA) in serum upon administration of etomoxir in uninfected mice. N=4-5 mice per group. The data are representative of two independent experiments. The figures show the mean ± s.e.m. Unpaired T test. \**p*<0.05, \*\**p*<0.005.

**Extended Data Fig. 9:**
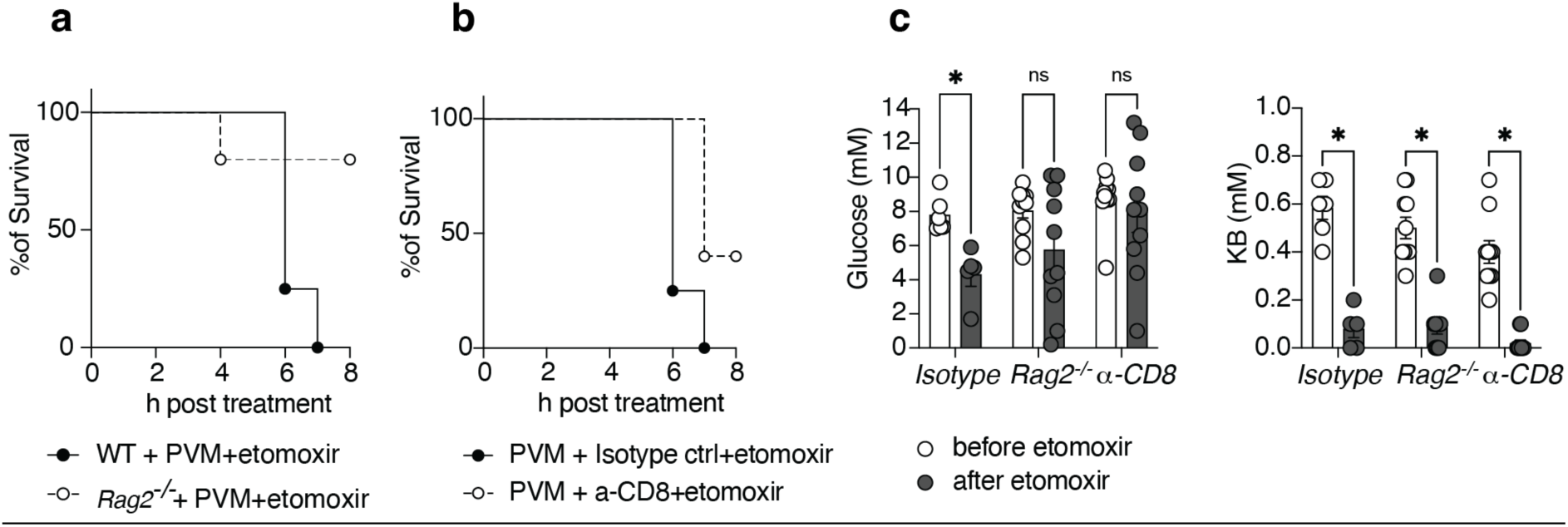
Characterization of PVM infected *Rag2*^-/-^ and CD8^+^ deficient mice. **a**, **b**, Nine days after infection with PVM, *Rag2^-/-^* mice or mice pretreated with a α-CD8 antibody, received a dose of etomoxir and survival was monitored every hour. N=4-5 mice per group, data is representative of two independent experiments. **c,** Blood glucose and KB were measured before and after etomoxir administration. N=5-10 mice per group, data pooled from two independent experiments. The data show the mean ± s.e.m. Unpaired T test. \**p*<0.05.

**Extended Data Fig. 10:**
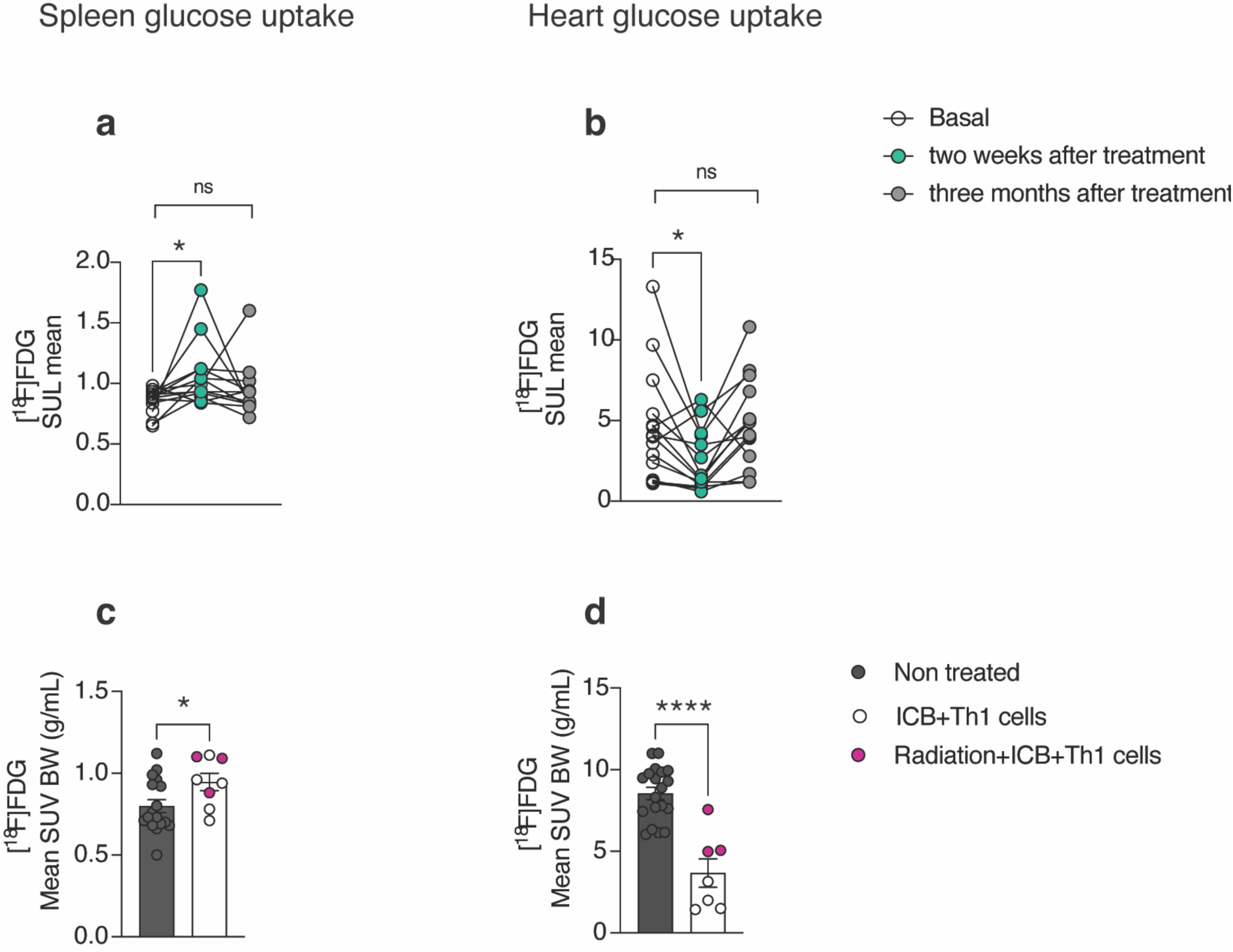
Glucose uptake in patients and mice receiving immunotherapy. Uptake of the [^18^F]FDG tracer was retrospectively analysed in the images obtained from patients with unresectable melanoma before treatment initiation (basal), two weeks and three months after initiation of immunotherapy. **a**, uptake of tracer in the spleen and **b**, heart expressed as standardized uptake value (SUV) mean corrected for lean body mass (SUL). N=14. Paired T test. **c**, **d**, Splenic and myocardial [^18^F]FDG uptake tracer was retrospectively analysed in the images obtained from mice with insular cell carcinoma that received no treatment or immune checkpoint blockade treatment (a-PDL1, a-LAG3) in combination with adoptively transferred tumor-specific Th1 cells, with our without prior low dose whole body radiation. Tracer uptake is expressed as SUV corrected for body weight. N=8-17. Data pooled from three independent experiments. The data show the mean ± s.e.m. Unpaired T test. \**p* <0.05, \*\*\*\**p* <0.0001.

